# Early functional mismatch between breast cancer cells and their tumour microenvironment suppresses long term growth

**DOI:** 10.1101/2021.06.15.448466

**Authors:** Anna Perdrix Rosell, Oscar Maiques, Probir Chakravarty, Luigi Ombrato, Victoria Sanz-Moreno, Ilaria Malanchi

## Abstract

Cancer cells thrive embedded in a fine-tuned cellular and extracellular environment or tumour microenvironment (TME). There is a general understanding of a co-evolution between cancer cells and their surrounding TME, pointing at a functional connection between cancer cells characteristics and the perturbations induced in their surrounding tissue. However, whether this functional connection needs to be set from the start or if aggressive cancer cells can always be dominating their microenvironment has never been formally proven with a dedicated experimental setting where malignant cells can be challenged to grow in a different TME from the one they would naturally create. Here we generated an experimental setting where we transiently perturb the secretory profile of aggressive breast cancer cells without affecting their intrinsic growth ability. This led to the initial establishment of an atypical TME. Interestingly, even if initially tumours are formed, this atypical TME evolves to impair long term *in vivo* cancer growth. Using a combination of in vivo transcriptomics, protein arrays and in vitro co-cultures, we found that the atypical TME culminates in the infiltration of macrophages with STAT1^high^ activity. These macrophages show strong anti-tumoural functions which reduce long-term tumour growth, despite lacking canonical M1 markers. Importantly, gene signatures of the mesenchymal compartment of the TME, as well as the anti-tumoural macrophages show striking prognostic power-correlating with less aggressive human breast cancers.

## Introduction

Cancer cells have the ability to generate a special cellular and extracellular environment to support their growth, which is collectively named tumour microenvironment (TME). This local ecosystem comprises an acellular part, characterized by a dense network of extracellular matrix (ECM) proteins, as well as a variety of cellular components (1). For instance, cancer-associated fibroblasts (CAFs) are one of the more dominant components of the TME and master secretors of ECM proteins and play a key role in tumour progression, tumour invasion and metastasis (2–4). Moreover, a multifaceted array of innate immune-cells, such as monocytes, macrophages and neutrophils, promotes tumorigenesis and metastatic dissemination (5–7). However, macrophages, which have a pivotal role in orchestrating cancer related inflammation, can have a dual supportive or inhibitory role in the cancer context (8). Importantly, the type of intra-tumoral immune infiltration in different cancer contexts is determined by the properties of the TME, for instance CAFs can contribute to the recruitment of immunosuppressive cells: tumour-associated macrophages (TAMs) or regulatory T lymphocytes (9).

An essential aspect of aggressive cancer cells is their ability to change their cytoskeleton to grow, invade, metastasize and evade the immune system. A key regulator of the cytoskeleton is Rho-associated coiled-coil containing protein kinase (ROCK) which increases Myosin II activity (10) and cancer invasion. Aside from its well described role in metastasis (11), ROCK has been shown to promote intrinsic tumorigenic ability of tumours of mesenchymal origin such as melanoma (12–14). Importantly, the tumorigenic potential of cancer cells is coupled to their extrinsic ability to promote a tumour supporting TME (15). Indeed, the fact that more contractile melanoma cells have a dinstinct secretory profile which alters the properties of TME components (1,16), advocate for the idea that the activation of pathways supporting cancer cells tumorigenic potential is directly linked to their ability to create their optimal cellular ecosystem within that tissue.

The fact that different tumour types have different cellular composition in their TME (17), rises the intriguing possibility that a specific match between the intrinsic characteristics of cancer cells and the kind of TME established is required to support their growth. In this view, it would be critical for a tumour to achieve a co-ordinated and fine-tuned TME establishment. In this case, a variation in the initial cancer cell-TME set up could result in an enduring mismatch that will ultimately impair tumour progression. However, whether changes in the initial TME typical of a particular cancer cell type, can act as dominant limiting factor for long-term tumour growth, or whether highly tumorigenic cells are able to re-adjust their TME in order to proficiently grow, is not known. This is due to challenging experimental set up required to address this question.

Here, we present an experimental setting where the secretion profile of the aggressive murine breast cancer 4T1 cells is perturbed by transitory inhibition of ROCK-Myosin II signalling either via the pre-treatment with a specific inhibitor or the use of inducible shRNA. Crucially, this does not influence 4T1 growth ability in vitro, but triggers a transient perturbation in the release of extrinsic factors that will initiate a different TME compared to the one they would normally create when grafted in the fat pad. Even when, at early stages of tumour growth, cancer cells recover their initial properties, we found that such TME limits their long-term cancer growth. This is due to the recruitment of a different stroma and the generation of anti-cancer macrophages. Therefore, this work represents a direct experimental evidence that the creation of an early optimal microenvironment is a dominant factor in the cascade of events determining long-lasting cancer growth and describe the signature of unusual anti-cancer macrophages which maintain M2 markers.

## Results

### Experimental uncoupling of cancer cells intrinsic growth potential and ability to activate stromal cells *in vitro*

To assess whether a variation in the initial cancer cell-TME set up could result in an enduring mismatch that will ultimately impair tumour progression, we sought a way to reversibly interfere exclusively with extrinsic features of cancer cells without negatively influencing their intrinsic tumorigenic potential. This should alter the initial interaction with their local environment and in turn generate a different TME. However, the cancer cells need to quicky regain their initial characteristics, to create the situation of a mismatch between the cancer cell type and its typical microenvironment *in vivo*.

Melanoma cells have been previously shown to have altered secretome upon treatment with the highly specific ROCK inhibitor GSK269962 (from now on named GSKi) (12,16). However, in melanoma, this signalling is generally essential for their growth potential. Conversely, human breast cancer lines show a variable level of intrinsic susceptibility to GSKi, and a substantial number of cell lines do not show an intrinsic response to the drug when treated *in vitro* for 3 days (Figure 1A). If alterations in the secretome of breast cancer cells is maintained, this treatment could potentially uncouple extrinsic features from intrinsic requirements. We therefore used GSKi to select a murine cell line able to maintain intrinsic growth capacity upon GSK addition and test if this still affected their secretion pattern. Cells were acutely exposed to GSKi for 5 days and their growth in different conditions was assessed after treatment interruption (pGSKi or pre-treated) (Figure 1B). The lower levels of MLC2 phosphorylation (pMLC2) confirmed that inhibition was maintained for at least three days after stopping the treatment in the murine 4T1 breast cancer cell line (Figure 1C and D). While pGSKi in other cell line, the 4T07 breast cancer line, caused reduced long-term growth (Supplementary Figure 1A and B), we found that did not reduced 4T1 cell growth in different *in vitro* conditions. (Figure 1E-G and Supplementary Figure 1C and D). Importantly, during this period, less contractile 4T1 cells showed an alteration in various secreted factors found in the culture media (Figure 1H and I). To assess the functional relevance of this perturbation in the secretome, we used classical fibroblast activation assays *in vitro*. After confirming the absence of GSKi in culture media of pre-treated cells by mass spectrometry (Supplementary Figure 1E), we found that both control and pGSKi-4T1 cancer cells similarly influence fibroblast proliferation *in vitro*. Nonetheless, these fibroblasts displayed different functional characteristics such as morphology, extracellular matrix contraction and migration abilities (Supplementary Figure 1F-J). These data indicate that, in principle pGSKi-4T1 cancer cells altered normal fibroblasts’ activation *in vitro*. As tumour associated fibroblasts are the most abundant and a key cellular component of the stroma in breast cancer (18), this data suggest that pre-treatment with GSKi could effectively uncouple intrinsic growth abilities and extrinsic host activating functions in 4T1 cancer cells, creating the ideal setting to address our research question.

**Figure 1.**
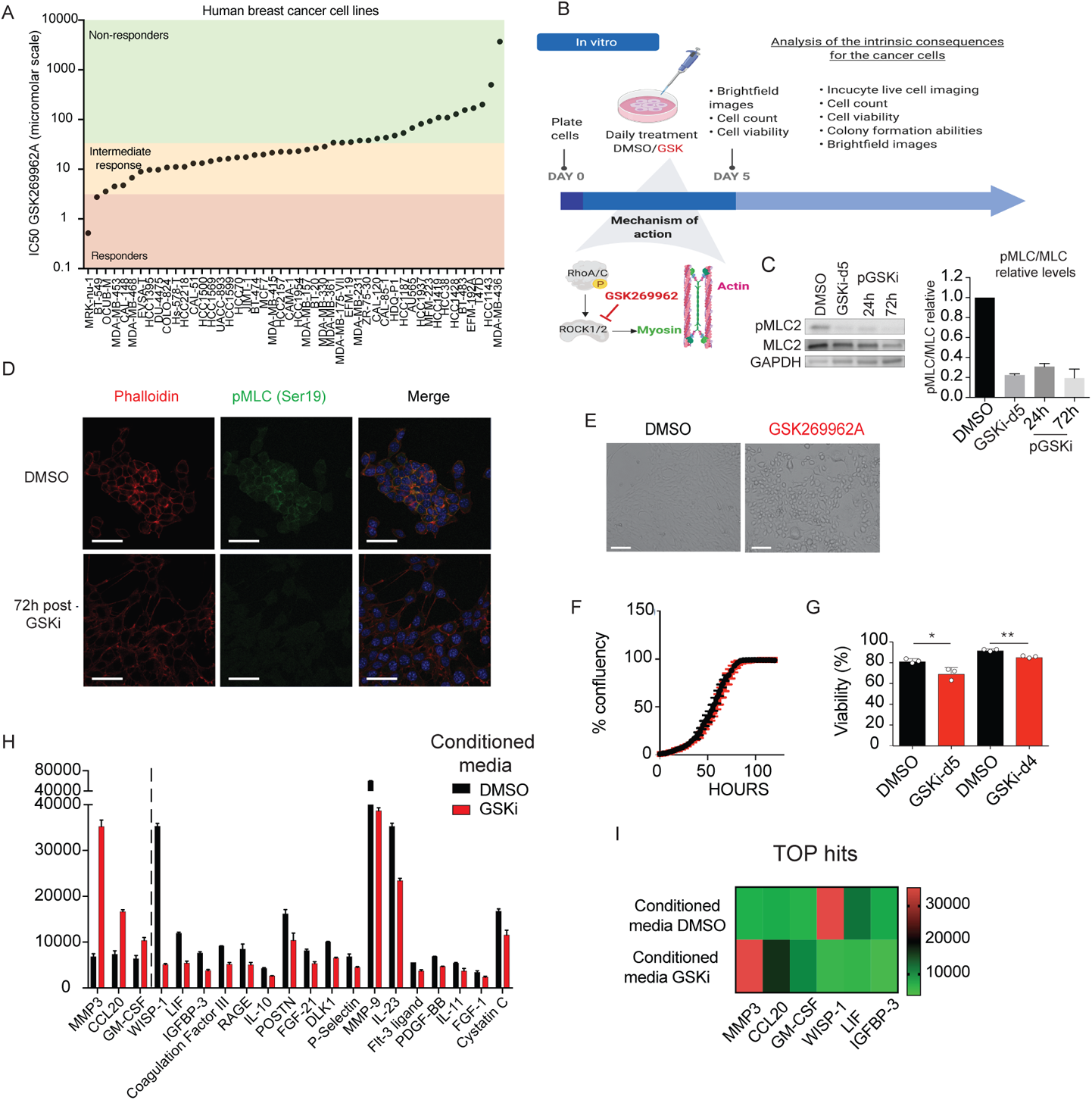
**A.** Distribution of GSKi response across human breast cancer cell lines. IC50 of GSK269962A inhibitor across 48 human breast cancer cell lines. Compounds are ranked in descending order of response and divided into responders (IC50<5μM, red), intermediate response (IC50 5-50μM, orange) and non-responders (IC50 >50μM, green). **B.** Schematic of the experimental setup for the characterisation of the intrinsic effects of GSK269962A on cancer cells. Cancer cells were plated and treated daily with vehicle (DMSO) or 5 μM of GSK269962A GSK1/2 inhibitor (GSK1) for five days. **C.** Western blot analysis of Phospho-MLC (pMLC) after 5 days of GSKi treatment (5 Days), 24h or 72h post-treatment. (Left panel) Quantifications of relative pMLC2 levels in 4T1 cells as a fold change versus the control from two independent experiments (right panel) **D.** Immunofluorescence images of pre-treated 4T1 cells 72 hours after treatment was removed. Cytoskeleton was stained using phalloidin (red), contractility using pMLC (green) and nucleai using DAPI (blue). (Scale bar 50μm). **E.** Representative brightfield images of 4T1 cells following a 5-day treatment with GSK269962A inhibitor. (Scale bar 100μm). **F.** Growth curve of cells pre-treated with DMSO (black) or GSK269962A inhibitor (red). Cell confluency was monitored every 3 hours for <120 hours using Incucyte. **G.** Percentage of cell viability quantified after 5 days of treatment and at day 4 post-treatment: DMSO (black) or GSK269962 (red) for 5 days. Error bars represent the SEM from three independent experiments. Statistic, Student’s t-test (*p < 0.05, ** p < 0.01). **H.** Quantification of differentially secreted factors in the conditioned media using Protein Array () of pre-treated 4T1 cells, 48 hours after treatment. Error bars represent SEM from two technical replicates. **I.** Heatmap of the top 3 upregulated (red) and downregulated (green) factors in GSK269962A-conditioned media.

### Early alterations in the TME causes long-term growth inhibition *in vivo*

Next, we tested the capacity of 4T1 cells pre-treated with GSKi (pGSKi-4T1) to form tumours when implanted in the fat pad of syngeneic Balb/c mice (Figure 2A). We found that while the initial phase of tumour establishment (8 days post-transplantation) was unaffected, the long-term growth ability of pGSKi-4T1 cells was strongly compromised (from 14 days post-transplantation) (Figure 2B and C). Importantly, at an early stage of tumour establishment-a time point where tumour growth was unaltered, the overall levels of pMLC2, monitoring the effect of the inhibitor, were nearly fully recovered from the GSKi pre-treatment, confirming its transient effect in vivo (Figure 2D and Supplementary Figure 2A-C). This unaltered early growth is in line with the unaffected intrinsic ability of cancer cell growth observed *in vitro* in pGSKi-4T1 (Figure 1E-G and Supplementary Figure1A and B). The long-term reduction of tumour growth resulting from GSK treatment was similarly achieved by specifically removing its target effector MYL9/MYL12B (genes encoding Myosin Light Chain 2) with an inducible shRNA (Supplementary Figure 2F-H). In this case, the effect in cancer cells was induced transiently as the mice were maintained under doxycycline induction only for 5 days post-transplantation. Similarly, to the results using GSKi pre-treatment, this resulted in an initial unperturbed growth (day 10), follow by a growth reduction and a long-term growth suppression (Supplementary Figure 2F-H).

**Figure 2.**
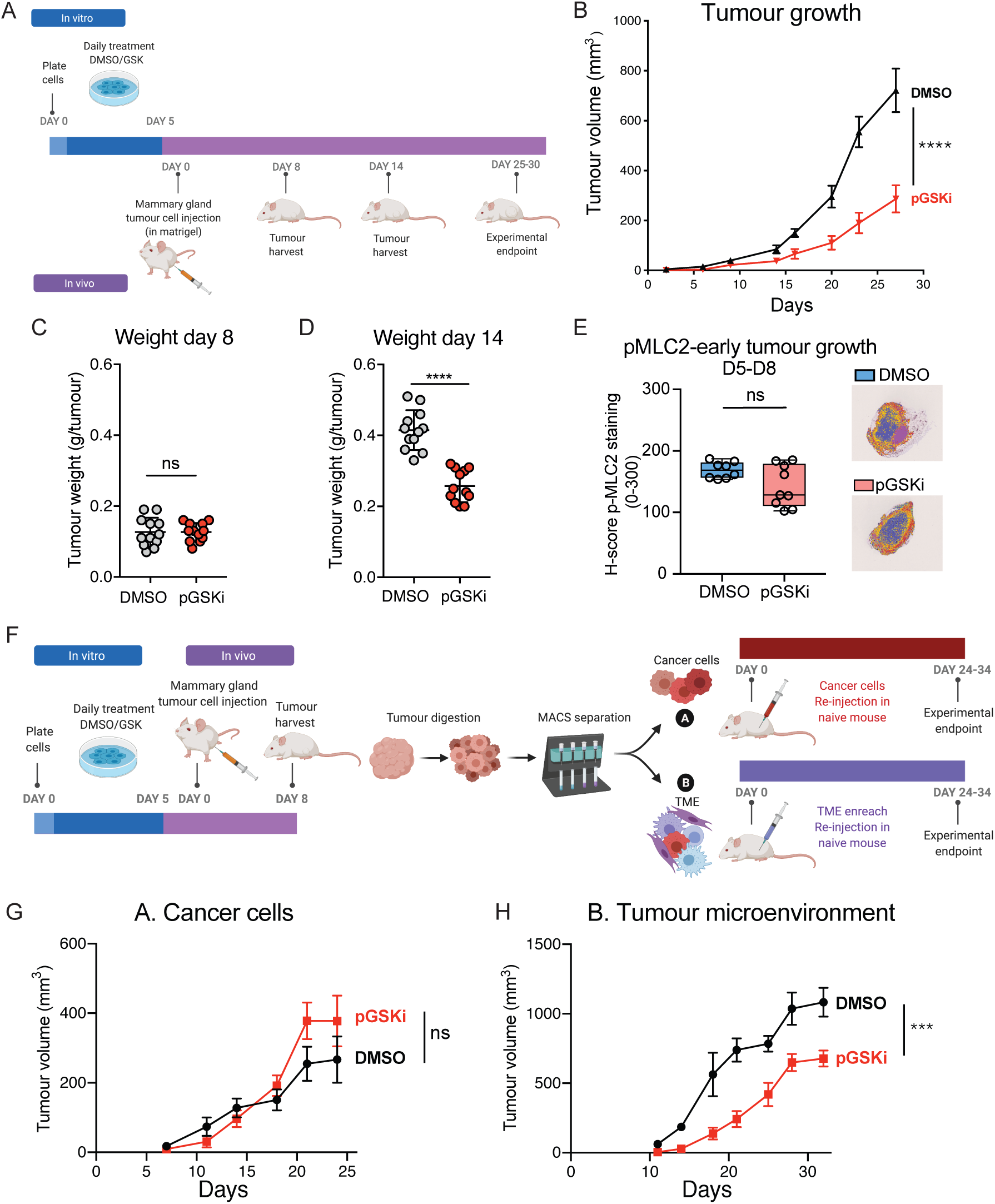
**A.** Schematic of the experimental setup. **B.** Mean tumour volume over time of tumours generated by 4T1 cells pre-treated with DMSO (black) or GSK269962A (red) in the fat pad of Balb/cAnN mice. Error bars represent SEM from four independent experiments (total of n=20 tumours). Statistic two-way ANOVA with Sidak’s multiple comparisons test **C-D.** Histogram of tumour weight at day 8 (n=28) (C) and day 14 (n=12) (D). Statistic: Student’s t-test between DMSO and GSK269962A groups. Statistically significant (ns: not significant, **** p < 0.0001 **E.** (Left) H-score of p-MLC2 staining in 4T1 tumour-derived cells pre-treated with DMSO (blue) or GSK269962A (red) from days 5 and 8 (Right) Representative QuPath color-maps of pMLC2 staining in both conditions. **F.** Schematic of the experimental setup. Day 8 tumours were separated by magnetic cell sorting into (A) cancer cells (hCD2+) and (B) tumour microenvironment (TME, hCD2-). Each of the components, was separately injected into new naïve Balb/cAnN mice. **G.** Mean tumour volume of (A) 4T1 cells isolated from day 8 DMSO (black) or GSK269962A (red) tumours. Error bars represent SEM from two independent experiments (n=9 total number of tumours). **H.** Mean tumour volume of (B) TME cells isolated from day 8 DMSO (black) or GSK269962A (red) tumours. Error bars represent SEM from two independent experiments (n=7 total number of tumours). Data were analysed using two-way ANOVA. (ns: not significant, *** p < 0.001).

Given that we have shown that pre-treated with GSKi altered the cancer-stromal interactions in vitro and that pGSKi-4T1 tumours suffer a delay in growth after the effect of the inhibitor has disappeared, we hypothesised that this delay could be due to the initial altered TME that might be less fit to support long term tumour growth. To test this hypothesis, we isolated cells from early-stage tumours, when tumour growth is not affected, and cancer cells have recovered from GSKi effects (Figure 2E). For this experiment we used 4T1 cells which were labelled by a extracellular domain of a human CD2 (hCD2) receptor. By magnetic cell sorting (MACS), we isolated either CD2+ cancer cells (A) or, alternatively, the rest of the TME depleted cells from the majority of cancer cells (B) (Figure 2F). By re-transplanting the isolated cells in new recipient mice, we tested the ability of 4T1 cells from pGSKi-4T1 tumours to re-establish secondary tumour growth alone or when maintained in the overwhelming presence of the TME they have previously established (Figure 2F). In line with the fact that GSKi effects was now cleared (Figure 2E and Supplementary Figure 2A-C), 4T1 cells from pGSKi-4T1 tumours show no delay in sustaining long term growth upon re-transplantation compared to control cells (Figure 2G). Strikingly, when the same cancer cells were re-transplanted together with the previously established TME, tumours still show a strong reduction of growth in secondary recipient (Figure 2H). These results point at a dominant inhibitory effect of the TME generated in pGSKi-4T1 tumours on cancer cells, which otherwise would possess an unaltered growth capacity.

### Early differences in stromal cells in inhibitory TME

We next analysed the TME cellular composition of tumours from control and pGSKi-4T1 cells after cancer establishment. Early pGSKi-4T1 tumours showed an increase in *α*SMA+ stromal cells, one of the dominant TME components already in control 4T1 tumours (Figure 3A and B). To test if the characteristics of the stromal compartment were different in the inhibitory TME, we isolated the general stromal compartment, depleted from immune-cells and endothelial cells, of tumours from pGSKi-4T1 cells (Figure 3C) at the early time point, when growth was unaltered. Deep RNA sequencing showed a different signature of stromal cells from pGSKi-4T1 compared to stromal cells isolated from control tumours (Figure 3D and E). However, when we functionally tested these alternatively activated stromal cells for their ability to support cancer cell growth, no difference was detected neither in 3D co-culture system nor in co-injection assay *in vivo* (Supplementary Figure 3A-D). We had previously observed that fibroblast activation *in vitro* was perturbed in pGSKi-4T1 cells (Supplementary Figure 1F-J) and similarly, we observed no difference in tumour growth also when co-injecting *in vitro* activated fibroblasts (Supplementary Figure 3E and F). This suggests that these differently activated stromal cells do not show anti-tumour effect per se, but they reflect the early generation of an altered TME initiated by pGSKi-4T1 cells.

**Figure 3.**
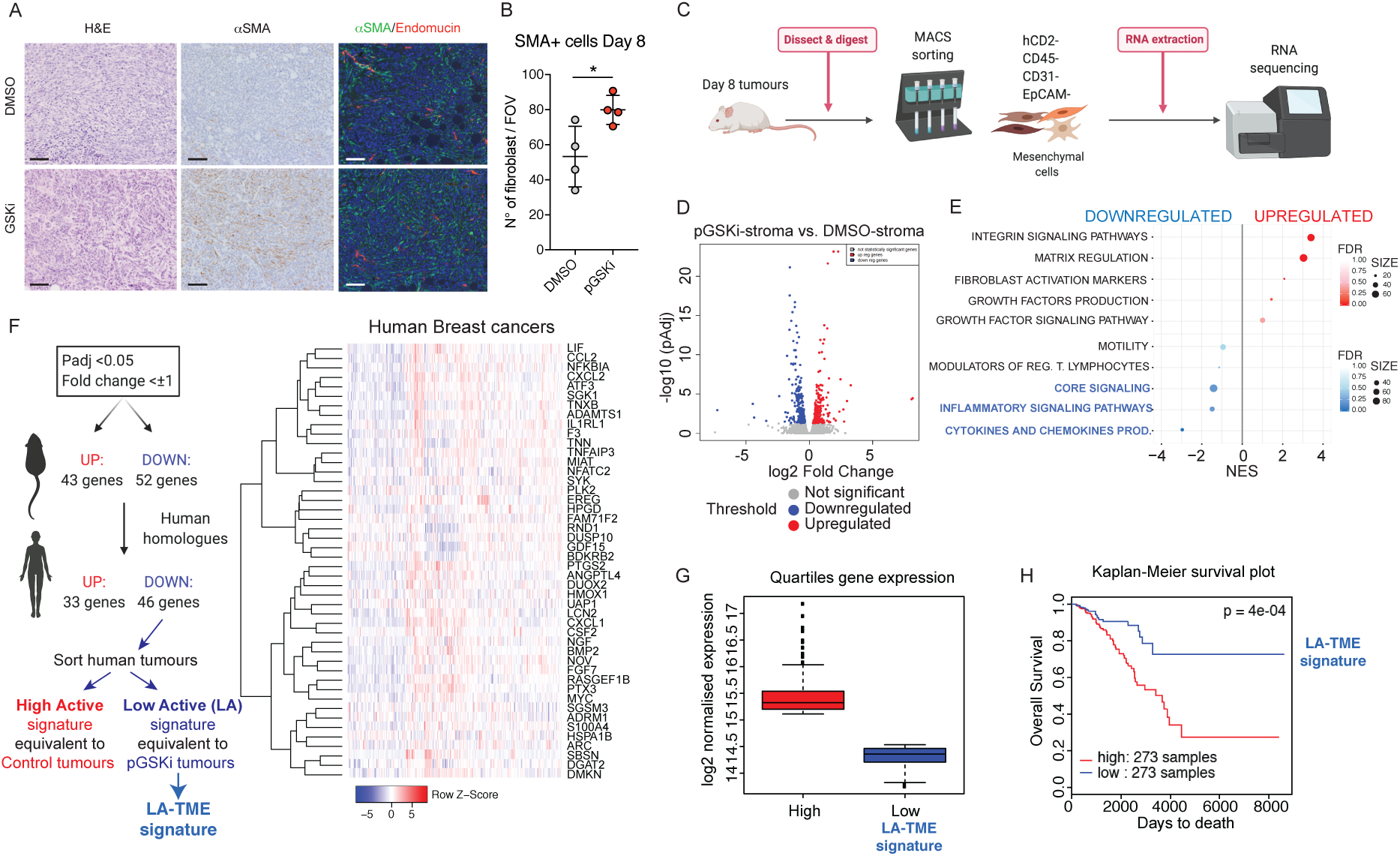
**A.** Representative IHC images of orthotopic syngeneic tumours 8 days post-transplantation from 4T1 pre-treated with DMSO or GSKi. Hematoxilin/Eosin (left), *α*SMA antibody DAB-staining (middle) and fluorescence double staining of *α*SMA (green), Endomucin (red) and DAPI (blue) to stain the nucleI (right) (Scale bar 100μm). **B.** Quantification of *α*SMA+Endomucin-cells in DMSO or GSKi pre-treated tumours 8 days post-transplantation. Error bars represent the SEM from 4 tumours with 6 fields of view (FOV)/tumour. Data were analysed using a student’s t-test (*p < 0.05). **C.** Experimental design for tumour associated mesenchymal cell isolation. **D.** Volcano plot comparing the gene expression profile of stromal cells derived from GSKi vs. DMSO pre-treated tumours. Statistical significance (-log10 adjusted pvalue) against foldchange (log2) of fibroblasts derived from GSKi vs. DMSO pre-treated tumours. Significantly upregulated genes are shown in red and downregulated genes are shown in blue. **E.** Gene signatures in CAF-specific molecular mechanisms were selected and ranked (upregulated in red; downregulated in blue). Gene-expression differences are displayed as a normalised enrichment score (NES), and false discovery rate (FDR) is used to determine the significance of each pathway. The size of the circle represents the number of genes within each signature that are differentially expressed in GSKi vs. DMSO-CAFs. **F.** (Left panel) Schematic representation of how the Cancer Low Contractility (CLC) - TME signature was generated stemming from genes downregulated in tumour initiated by cancel cells with reduced contractility (GSKi). Padjusted value (Padj) and log2 fold change (fold change) were used as cut-offs for gene selection. (Right panel) Heatmap of the 46 genes that constitute the CLC-TME signature and their expression across 1092 breast cancer tumours from the TCGA cohort. **G.** Log2 expression of the CLC-TME signature divided patients into high (red, equivalent to signature in mouse control tumours) and low (blue, equivalent to signature in mouse GSKi tumours) quartiles. Box-an-whisker plots showing median centre line, 25% and 75% box limits and rang of expression. **H.** Kaplan-Meier curve represents the overall survival of 546 breast cancer patients from the TCGA cohort subdivided into high (red) and low (blue) expression of the GSKi-CAF “down signature”. Data were analysed using a Wald test and statistically significant changes between the two groups are indicated (* p < 0.05) (hazard ratio is 1.96 and confidence intervals are 1.1 – 3.93).

When analysing the features of the alternative TME captured by its stromal signature, we found that extracellular matrix (ECM) regulation and integrin signalling were upregulated (Figure 3E) pointing at changes within the non-cellular ECM compartments of the TME. We could indeed validate this observation by detecting an increase in matrix density and Collagen IV staining in the pGSKi-4T1 tumours compared to control (Supplementary Figure 3G and H). Importantly, the stromal signature found in tumours generated by pGSKi-4T1 cells which show reduced activation, Low Active (LA)-TME signature, showed strong prognostic power to identify human breast cancer patients with better survival in TCGA database (Figure 3F-H). This indicates that this lower level of typically activated pathways in the stromal signature characterized less aggressive human breast cancers. Indeed, this LA-TME signature shows downregulation of pathways key for CAF function (Figure 3E), suggestive of a failure to establish the suitable pro-tumoral activity needed in the TME of aggressive and fast growing tumours.

### Inhibitory TMEs contain different immuno-infiltrates

Bioinformatic analysis showed that a major difference in the enriched pathways in pGSKi-4T1 tumour’s stroma was lower activation of inflammatory signalling pathways/chemokines and cytokine release, suggesting that broader alterations could be triggered within the inflammatory TME compartment (Figure 3E). Consistently, when the immune-cell composition of tumours was analysed by FACS, we found significant differences in control or pGSKi-4T1 tumours specifically at later time points (Figure 4A-H). Data showed a decrease in cells from the lymphoid lineage in favour of an increase in myeloid cells.

**Figure 4.**
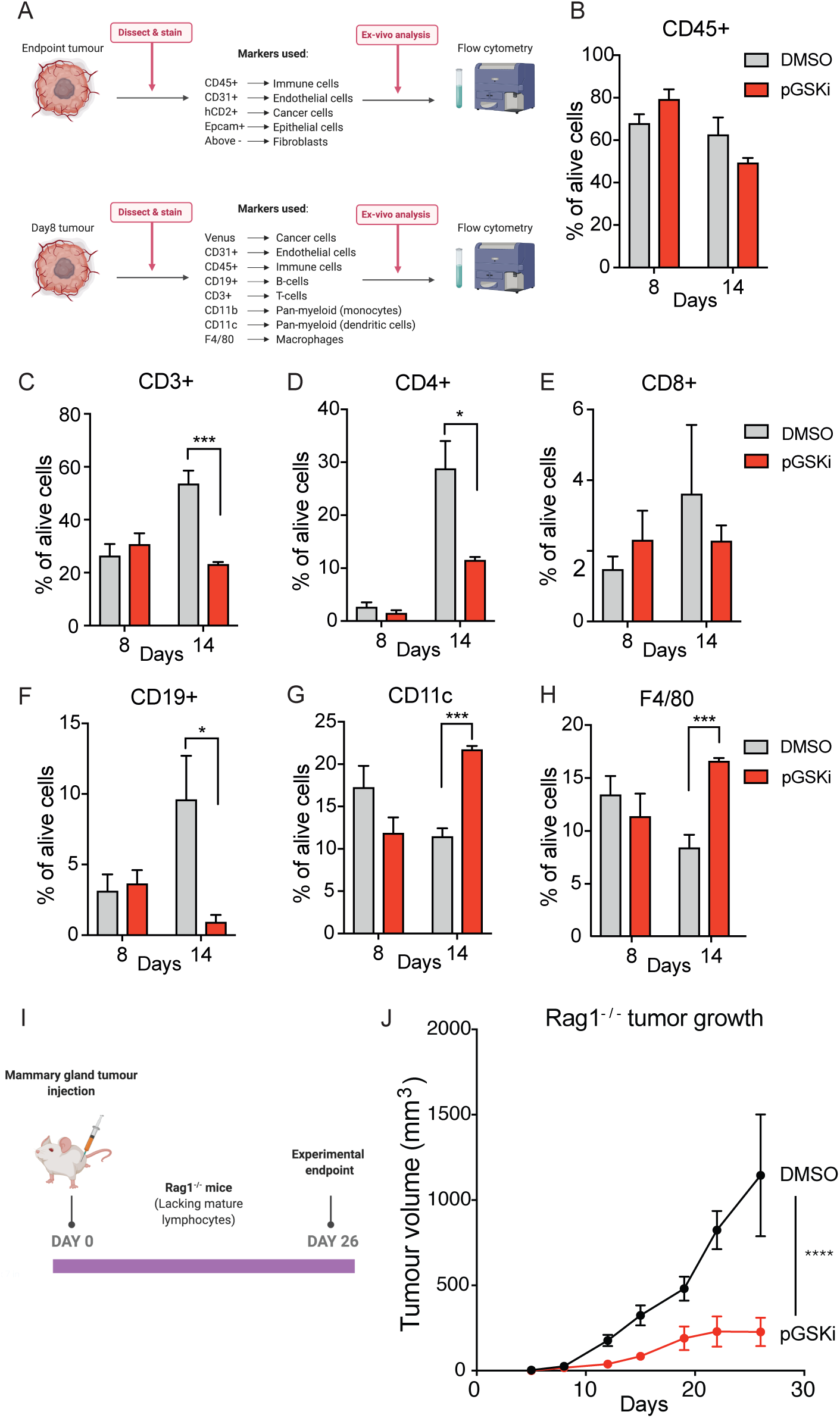
**A.** Experimental design and antibodies used for detecting immune cells populations within the TME. **B-H.** Flow cytometric quantification of TME cell frequencies out of the total alive cells lungs in mice harbouring DMSO (grey) or GSK269962A (GSKi) (red) pre-treated tumours at the indicated times post-injection. (B) total CD45+ immune cells (C) CD45+CD3+ T-cells; (D) CD45+CD4+ T-cells; (E) CD45+CD8+ T-cells; (F) CD45+CD19+ B-cells; (G) CD45+CD11c+ dendritic cells; (H) CD11b+F4/80+ macrophages. Error bars represent the SEM from two independent experiments (Total n=12 in each group). Data were analysed using a t-test between DMSO and GSKi groups at each time points. (* p < 0.05, *** p < 0.001). **I.** Diagram of the experimental design. **J.** Mean tumour volume of 4T1 cells grown over 26 days. Tumours were pre-treated in vitro with DMSO (black) or GSKi (red). Error bars represent the SEM from two independent experiments (total of n=12 (DMSO) and n=10 (GSKi)). Data were analysed using a two-way ANOVA (**** p < 0.0001).

To test if a decrease in tumour infiltrating lymphocytes could contribute to the delay in tumour growth, control or pGSKi-4T1 cells were transplanted in immune-compromised RAG1-/- mice, lacking B and T lymphocytes. Similar delays in the growth of pGSKi-4T1 tumours were observed, excluding an involvement of the adaptive immune system to limit tumour growth (Figure 4I and J).

Macrophages are known to possess both pro- and anti-tumorigenic functions (19), therefore we tested if different macrophage polarizationcould be detected in pGSKi-4T1 tumours. Tumour associated macrophages (TAM) showed no difference in canonical markers that have been previously associated to pro- or anti-cancer activities (Supplementary Figure 4A and B). Accordingly, when analysing the immune-infiltrates in the TCGA human breast cancers classified according to the pGSKi-4T1 tumours stromal signature, there was no difference in M1 or M2 macrophages (Supplementary Figure 4C). Collectively, these data show that the change in lymphocyte infiltration status of the pGSKi-4T1 TME it is not responsible for the reduced growth. Surprisingly, the macrophages infiltrating these tumours still express classical pro-tumorigenic markers.

### Inhibitory TMEs harbour macrophages with non-canonical anti-cancer activity

We next aimed to determine if, despite showing canonical TAM markers, macrophages in pGSKi-4T1 tumours, harbour different characteristics. To do this, we generated a gene expression signature of F4/80 macrophages isolated from control or pGSKi-4T1 tumours (Figure 5A). Striking differences were detected at the gene expression level (Figure 5B and C). Interestingly, the emergence of a distinct signature was detected 8 days post-transplantation and was consolidated at later time points, when tumour growth was significantly reduced (Figure 5B and C).

**Figure 5.**
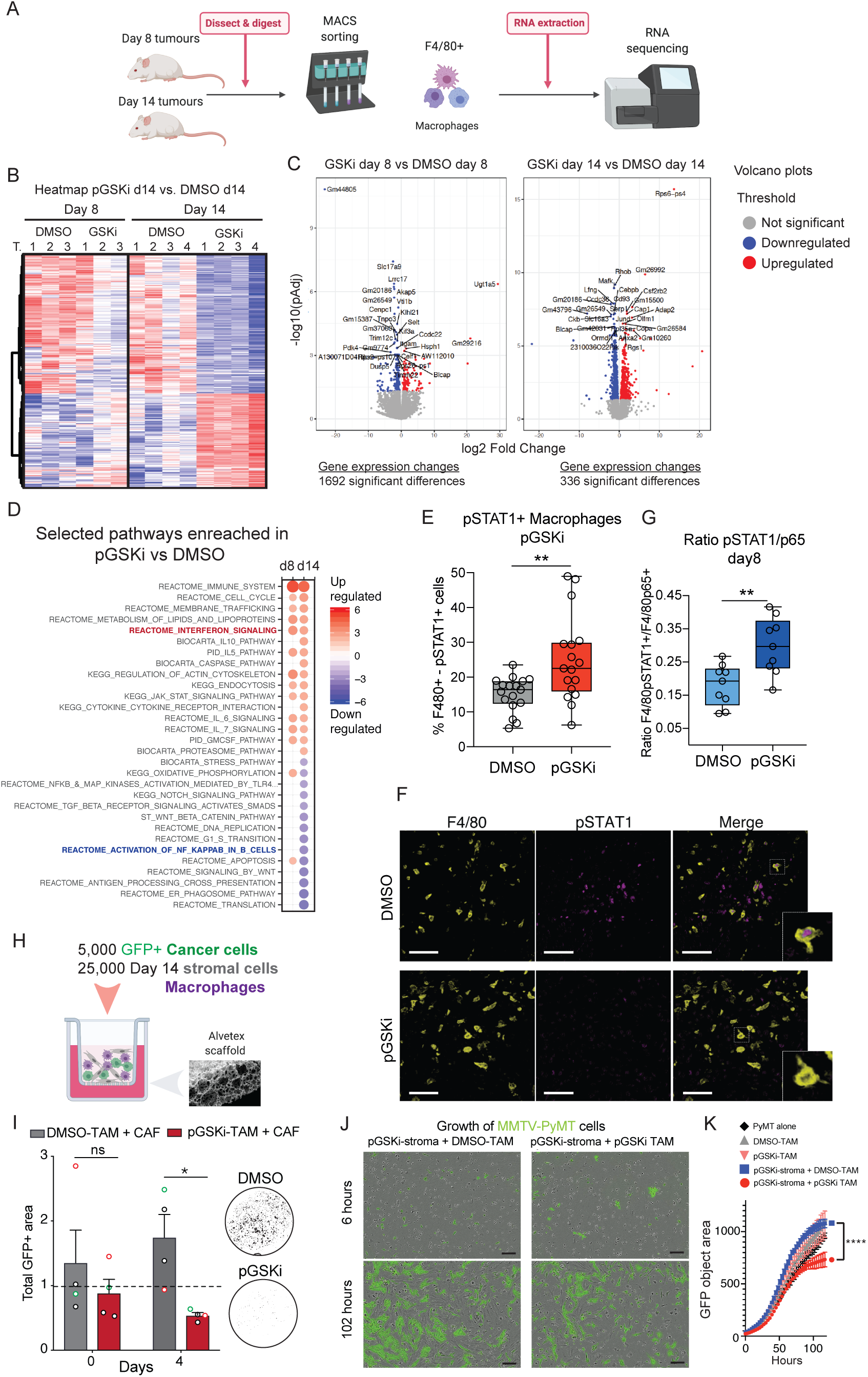
**A.** Schematic representation of the experimental design and sample collection. **B.** Heatmap comparing the gene expression profile of macrophages derived from GSKi vs. DMSO pre-treated tumours 8 and 14 days post-transplantation. **C.** Volcano plots of statistical significance (-log10 adjusted pvalue) against foldchange (log2) of macrophages derived from GSKi vs. DMSO day 8 pre-treated tumours (Left) or day 14 pre-treated tumours (Right). Significantly upregulated genes are shown in red and downregulated genes are shown in blue. **D.** Summary of selected pathways enriched in GSKi vs. DMSO. Upregulated or downregulated pathways at the indicated time point are indicated in in red and blue respectively. **E.** Percentage of pSTAT1 positive macrophages derived from DMSO (grey) or GSKi (red) pretreated tumours from day 14. **F.** Representative staining of F4/80 (yellow) and pSTAT1 (Magenta) of DMSO or GSKi pre-treated tumours section (Scale bar 50μm). **G**. Ration between p65 and pSTAT1 staining in macrophages in tumour pre-treated with DMSO (gray) or GSKi (red) from day 8. **H.** Experimental design of the co-culture. **I.** Quantification of primary MMTV-PyMT-GFP+ cell growth by measuring the total GFP+ area over the course of 4 days. Cells were grown with DMSO-Macrophages and CAFs (grey) or GSKi-macrophages and CAFs (red) from 14-days tumours and normalised to the growth of MMTV-PyMT cells alone (dash line). Representative fluorescence pictures are shown. Error bars represent the SEM from three independent experiments shown with black, green or red data points. Biological replicates within each experiment are indicated in the graph. Data were analysed using a Student’s t-test (* p < 0.05). **J.** Representative brightfield images of MMTV-PyMT-GFP+ cells (green) in co-culture with GSKi-CAFs and either DMSO (left) or GSKi (right) macrophages. Timepoints are indicated. Scale bar, 30 µm. **K.** Quantification of MMTV-PyMT-GFP+ cell area when cells were grown alone (black), with macrophages from DMSO (grey) or GSKi (coral) pre-treated tumours or with CAFs from GSKi pre-treated tumours and macrophages extracted from either DMSO (blue) or GSKi (red) pre-treated tumours. Statistic two-way ANOVA (blue vs red) (**** p < 0.0001).

Analysis of the macrophage signature from the two types of tumours using GSEA revealed an increase in anti-tumour inflammation Interferon gamma (INF*γ*) signalling and a decrease in pro-tumour inflammation NF*κ*B signalling in pGSKi-4T1 tumour derived macrophages (Figure 5D), pointing to an anti-tumour function. Importantly, when we analysed the conditioned media of stromal cells isolated from pGSKi-4T1 tumours at 14 days, we detected a dramatic decrease in specific secreted factors cytokines compared to control tumours (Supplementary Figure 5A). Particularly, at this tumour stage, network enrichment analysis of the secretome supported the overall decrease in NF*κ*B signalling within stromal cells (Supplementary Figure 5B), which was previously reported to be critical for CAF-mediated enhancement of tumour promoting inflammation (20). The activation of INF*γ* activity indicated by the signature of infiltrating macrophages of pGSKi-4T1 tumours, was confirmed by a sharp increase in active STAT1 nuclear staining in such macrophages at late time points (Figure 5E and F). Consistently with a growth defect , similar pSTAT1+ macrophages were detected at the late stage of tumours initiated by cells transiently affected by MYL9/12b knock down (Supplementary Figure 2D-F and 5E). Interestingly, already at early stages, infiltrating macrophages of pGSKi-4T1 tumours sowed increased pSTAT1 staining (Supplementary Figure 5C). Moreover, as predicted by their signature, we also measured a reduction in nuclear p65 (RelA component of the NF*κ*B complex) in macrophages from pGSKi-4T1 tumours (Supplementary Figure 5D), which results in an overall switch from NF*κ*B to INF*γ* -STAT1 signalling activity (Figure 5G). We therefore hypothesise that macrophages in pGSKi-4T1 tumours undergo profound transcriptional re-programming that might compromise their pro-tumorigenic potential without changing some key cell surface markers.

To assess the functional implications of the changes observed in macrophages, we tested their direct interaction with cancer cells by using a 3D co-culture scaffold system. We used primary breast tumour cells isolated from spontaneous Actin-GFP/MMTV-PyMT breast tumours in the co-culture. Freshly isolated macrophages from either control or pGSKi-4T1 tumours showed no differential activity when co-cultured alone with cancer cells seeded on 3D scaffolds (Supplementary Figure 5F-H). However, when cancer cells were seeded together with their cancer associated stromal cells, macrophages isolated from 14 days pGSKi-4T1 tumours showed a cancer inhibitory effect (Figure 5H and I). Similar results were observed in 2D culture conditions (Figure 5J, 5K). Fibroblasts supported the cancer cell killing activity of macrophages isolated from pGSKi-4T1 tumours independently from their origin. when isolated from pGSKi-4T1 tumours or control (DMSO) tumours (Figure 5K). These data suggest that the anti-tumour polarization observed in macrophages from pGSKi-4T1 is progressively established within the complex *in vivo* TME and it becomes a hardwired status.

### Non-canonical pGSKi-4T1 macrophages are responsible for limiting tumour growth

To test whether this non-canonical anti-tumour polarization of macrophages was acting to limit cancer cell growth *in vivo*, we used clodronate to deplete macrophages from control or pGSKi-4T1 cell transplants (Figure 6A). In line with the known pro-tumour activity of tumour infiltrating macrophages in breast cancer (19), their depletion decreased the final size reached by control tumours (Figure 6B). However, an opposite effect was shown in pGSKi-4T1, where their depletion boosted overall tumour growth and rescued it to the level of control tumours (Figure 6C). This data highlighted that the alternative macrophage polarization we described, indeed results in a switch from pro- to anti-tumour function.

**Figure 6.**
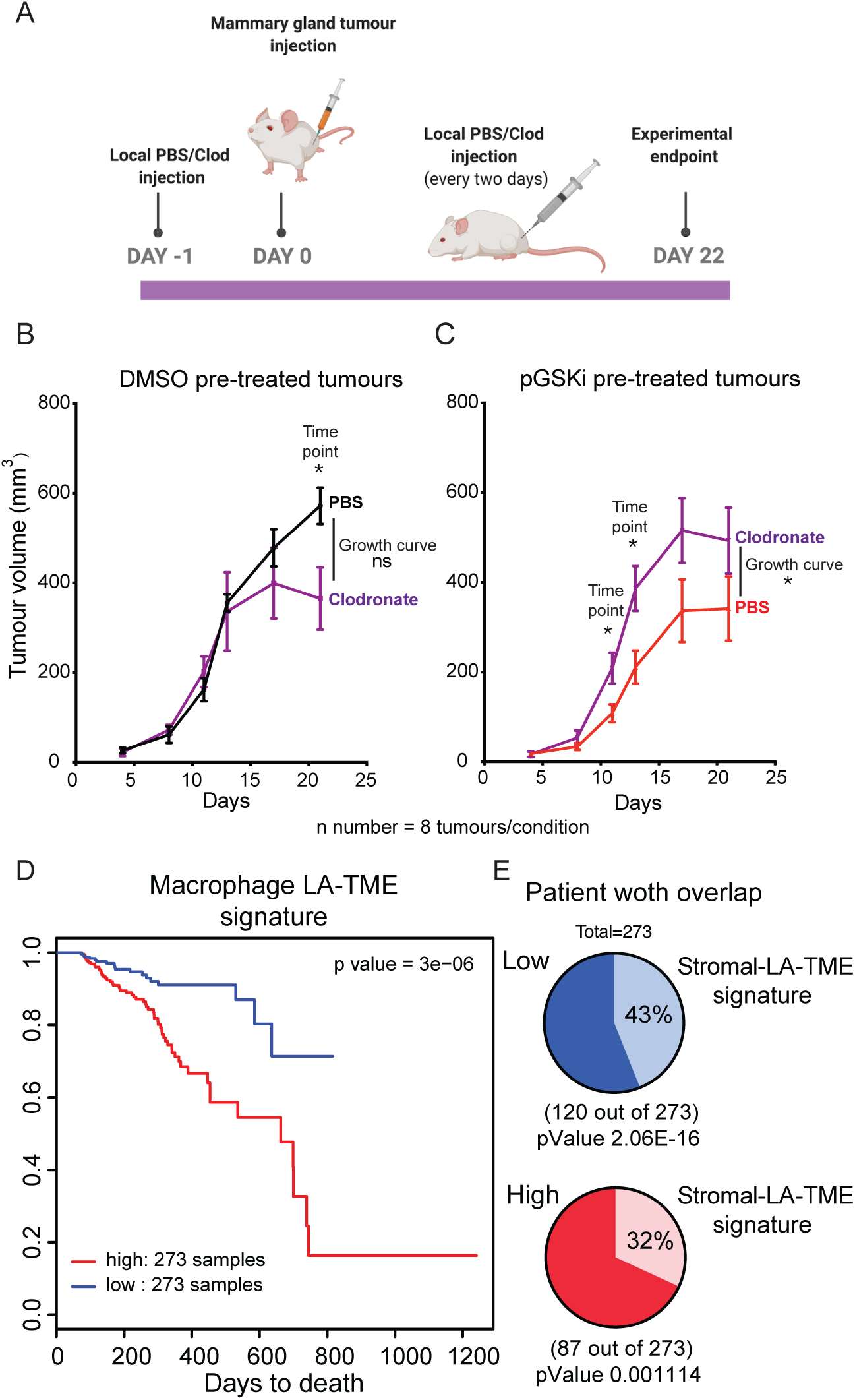
**A.** Experimental design. **B-C.** Mean volume of 4T1 tumours over 22 days. 4T1 tumour cells were pre-treated in vitro with DMSO (B) or GSK inhibitor (C), and tumours were treated with either PBS liposomes (black and red lines) or clodronate liposomes (purple lines). Error bars represent the SEM (n=8 tumours per condition). Data were analysed using a two-way ANOVA comparing the curves (at the end of the curves) (ns: not significant, * p < 0.05) and t-tests at individual timepoints when significant difference were present (indicated above each time point) (* p < 0.05). **D.** Kaplan-Meier curve represents the overall survival of 546 breast cancer patients from the TCGA cohort subdivided into high (red) and low (blue) expression of the macrophage signature. Data were analysed using a Wald test and statistically significant changes between the two groups are indicated (* p < 0.05) (hazard ratio is 1.96 and confidence intervals are 1.1 – 3.93). **E.** Pie charts represent the overlap between patients with both alternative macrophage and alternative stromal signatures from GSKi pre-ptreated tumours.

We have shown in Figure 3 that the activated stroma from pGSKi-4T1 cells harbour a gene signature (LA-TME signature) with prognostic value in human breast cancer. Similarly, we generated the macrophages LA-TME signature using the downregulated genes in macrophages from pGSKi-4T1 tumours. Strikingly, we found that the macrophage LA-TME signature showed a similar power in predicting better survival of human breast cancer patients (Figure 6D). Moreover, 43% of these human tumours additionally showed the alternative stroma LA-TME signature, suggesting a similar overall reduction of activation in stromal cells and macrophages genes in the tumour microenvironment can be found in human breast cancers with better prognosis (Figure 6E). Most importantly, as no difference was found in macrophage markers generally associated to anti-tumour phenotypes in breast tumours (Supplementary Figure 4A-C), this data suggests the presence of a previously unappreciated alternative type of anti-tumour macrophages, which correlates with better clinical outcome.

Overall, this study show that an initial deviation in the initial stage of cancer-host interaction cause by a perturbation of the malignant cancer cells secretion, results in the long term establishment of a suboptimal TME which becomes dominant in suppressing aggressive tumour growth.

## Discussion

In this work, we show that the initial TME typical of a particular cancer cell type, can act as dominant limiting factor for long term tumour growth. To test this, we have generated an experimental set-up which allows to interfere exclusively with the extrinsic features of an aggressive cell line without negatively influencing its intrinsic growth potential. This change induces an initial atypical interaction with the *in vivo* environment which is different from what these cancer cells would induce. In turn, this generates an initial alternative TME. As the nature of the perturbation induced in the cancer cells is temporary, they recover their initial characteristics when embedded in this atypical microenvironment *in vivo.* By creating this complex experimental strategy, we here formally show that a sub-optimal TME becomes dominant in suppressing the growth of highly aggressive cancer cells. In this work, also show that ROCK-Myosin II activation in this breast cancer cell line with basal characteristics control their secretion activity and modulate the initial activation of host cells in vitro and in vivo without altering their intrinsic growth potential. This is intriguing as cancer cells displaying high ROCK-Myosin II activation, generally relay on this pathway for their intrinsic tumourigenicity (16), but we identify a murine breast cancer cell line which represented an exception and where it exclusively interferes with the secretion profile of the cells. Indeed, the aggressive murine 4T1 breast cancer maintained its highly proliferative activity in various vitro conditions upon ROCK-Myosin II inhibition but showed a perturbed fibroblast activation ability. After identifying the 4T1 as a cell line where the intrinsic and extrinsic activity could be uncoupled using ROCK-Myosin II inhibition, we used ROCK-signalling inhibitor (GSK269962A) (GSKi) pre-treatment before transplanting cancer cells *in vivo* to only transiently perturbe their interactions with the host environment. Even when the GSKi effect ceased and cancer cells recover ROCK-Myosin II activity, are now less capable to sustained long-term growth. Secondary transplantation experiments show that, while cancer cells recover their full in vivo tumourigenicity when growing secondary tumours, is the TME generated by initially pre-treated GSKi 4T1 cells who imposes a limitation for their growth.

Our data supports the fact that GSKi pre-treated cancer cells initiate an atypical TME characterized by an increase early representation of SMA positive cells, which, similar to the in vitro activated fibroblasts, show different characteristics and signature compare to the one that unperturbed 4T1 would have generated. Secretome analysis on GSKi pre-treated 4T1 conditioned media revealed significant differences with control 4T1 media in the secretion of 20 cytokines (where LIF, WISP1, MMP9 and several others resulted downregulated in pGSKi), some of which have been previously linked to CAF activation. For instance, cancer-derived LIF was shown to promote CAF activation and pro-invasive phenotype via inducing actomyosin contractility and ECM remodelling (21). In addition, WISP1 was found in the metastatic niche and was required for breast cancer metastasis to the lung (22). Thus, it is tempting to speculate that such a differential cocktail of factors could interfere with the pro-tumorigenic stromal activation generally induced by 4T1 cells.

Strikingly, despite the transcriptomic analysis of stromal TME component from control or GSKi pre-treated tumours shows an atypical gene-expression signature, they similarly supported cancer cell growth *ex-vivo* and *in vivo*. This data suggests that in principle, they can support cancer cell growth. Nonetheless, within the TME, this atypical stromal component is part of a more complex environment, which ultimately results to be unfit to support aggressive tumour growth. In particularly, we detected a reduced activation of inflammatory signalling NFkB, which iscrucial for the establishment of pro-tumoral inflammation (20). This lack of activation has the potential to have a broader impact on the immune compartment in the TME. Indeed, when analysing the tumour infiltrates, we identified a reduction on lymphocytic cells (B and T-cells, specifically CD4+ but not CD8+ cells) and an increase in myeloid cells (macrophages and dendritic cells). We excluded the role of the lymphocytic compartment in the observed phenotype as GSKi pre-treated cancer cells maintained their growth defect in genetically-immunodeficient mice lacking mature lymphocytes (Rag1-/-).

Although TAMs did not present distinct expression of canonical pro- or anti-tumour markers between tumours from control and pGSKi cancer cells, RNAseq analysis showed broad changes in their transcriptome. In particular, TAMs from tumours generated by GSKi pre-treated cancer cells suggested an anti-tumour function with a significant increase in Interferon *γ* (INF*γ*) signallling. Indeed, TAM infiltrating pGSKi tumours showed clear increase in nuclear pSTAT1, the effector transcription factor downstream of INF*γ* . We could confirm that this anti-tumorigenic function of TAM was responsible for inhibiting cancer growth in *ex-vivo* and *in vivo* experiments. Interestingly, in *ex-vivo* co-culture, the TAM anti-tumour function was only detected in presence of freshly isolated stromal cells, but this did not depend on their origin, could either be from pGSKi or control tumours. This suggests that fibroblasts might enhance TAM toxicity by providing adhesion for macrophages and increase their proximity to cancer cells and also suggests that macrophage polarization acquired intra-tumorally is a stable phenotype independent from external factors.

Finally, when testing the functional role of macrophages on tumour growth *in vivo* we confirmed their anti-tumour function. By using Clodronate, a type of bisphosphonate which once encapsulated into liposomes is up taken by macrophages and mediates their depletion (23), we could clearly see the opposing effects in control or pGSKi tumours in either inhibiting or promoting growth respectively.

Thus far, it has been shown how CAFs can contribute to assembling an immunosuppressive TME accordingly to their type of activation (9). We could detect an alteration in pathways responsible with immunomodulation in the TME mesenchymal compartment. We found macrophages infiltrating pGSKi tumours to display increased INF*γ*-STAT1activation and reduced NFkB signalling. This was shown by the increase in nuclear staining for the INF*γ* effector pSTAT1 and reduced staining for p65. INF*γ* signalling is typical of anti-tumour functions in the immune-microenvironment (24), therefore a switch in signalling activation could control macrophage behaviour.

Overall, our data shows that cancer cells assemble very specific tumour microenvironments very early in tumour development. The present study reveals a key role of an atypical TME as dominant in inhibiting tumour growth due to the lack of crucial pro-tumoral signals leading to the onset of anti-tumour TAMs (Figure 7). The different TME contains fibroblasts and macrophages with Low Activation of clusters of genes, LA-TME signatures, which allows identification of human breast cancers with better prognosis.

**Figure 7.**
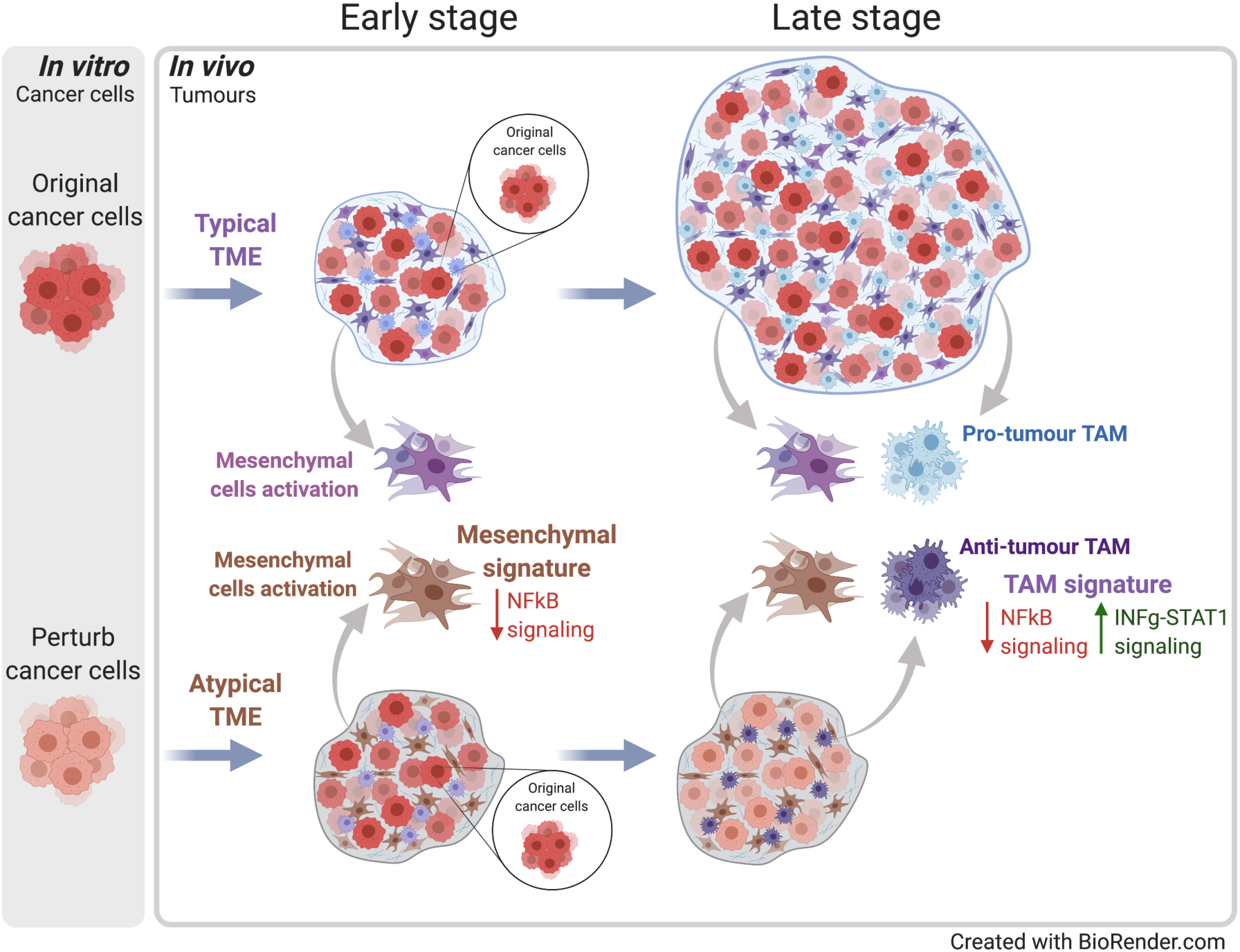
Graphic summary: breast cancer cells with initial reduced GSK-Myosin II activity generated an alternative early TME characterized by alternative Carcinoma Activated Fibroblasts (CAFs) displaying a Cancer Low Contractility (CLC)-CAF-signature. The CLC-CAF-signature show overall reduced pro-inflammatory activity including down-regulation of NF*κ*B signalling. The alternative TMA fails to sustain pro-tumourigenic signalling typical of Tumour Associated Macrophages (TAM), which consequently display a CLC-TAM-signature with reduced pro-tumoural NF*κ*B signalling and anti-tumour Interferon Gamma (INF*γ*), STAT1^high^ activity. These macrophages -lacking canonical M1 markers-reduced long-term tumour growth.

Remarkably, the TAMs found in tumours from GSKi pre-treated cells, despite displaying cancer killing activity, do not show canonical M1 markers nor M1 macrophage infiltration was observed in human tumours correlating with the LA-TME signatures. This suggesting that this non-canonical anti-tumour macrophage polarisation might exist in human tumours and contribute to better clinical outcome.

## Acknowledgments

The graphic representation in the figures were created using BioRender.com.

We thank E. Sahai (The Francis Crick Institute) for scientific discussions. We thank Jose L. Orgaz-Bueno, Eva Crosas-Molist, Irene Rodriguez-Hernandez, Stefania di Blasio, Victoria Bridgeman and Emma Nolan for scientific discussions and support during the course of this project. E. Nye from the Experimental Histopathology Unit at the Francis Crick Institute for histological processing and analysis support; J. Bee from the Biological Resources Unit at the Francis Crick Institute for technical support with mice and mouse tissues; R. Goldstone and A. Edwards from the Advanced Sequencing Facility at the Francis Crick Institute for technical support; the Flow Cytometry Unit at the Francis Crick Institute.

This work was supported by the Francis Crick Institute, which receives its core funding from Cancer Research UK (FC001112), the UK Medical Research Council (FC001112), and the Wellcome Trust (FC001112) and the European Research Council grant (ERC CoG-H2020-725492). VSM lab was supported by Cancer Research UK (CRUK) C33043/A24478 and Barts Charity.

## Author contributions

A.P.R. designed and performed most of the experiments, analysed and interpreted the data and contributed to the manuscript preparation. O.M. performed tumour staining and quantification analysis of in vivo macrophages, performed secretome data analysis and critically review the manuscript. L.O. assisted with some of the experimental design and critically review the manuscript. P.C. performed bioinformatics analysis. I.M. and V.S.M. designed and supervised the study, interpreted the data and wrote the manuscript.

## Supplementary Figures and Legends

**Supplementary Figure 1.**
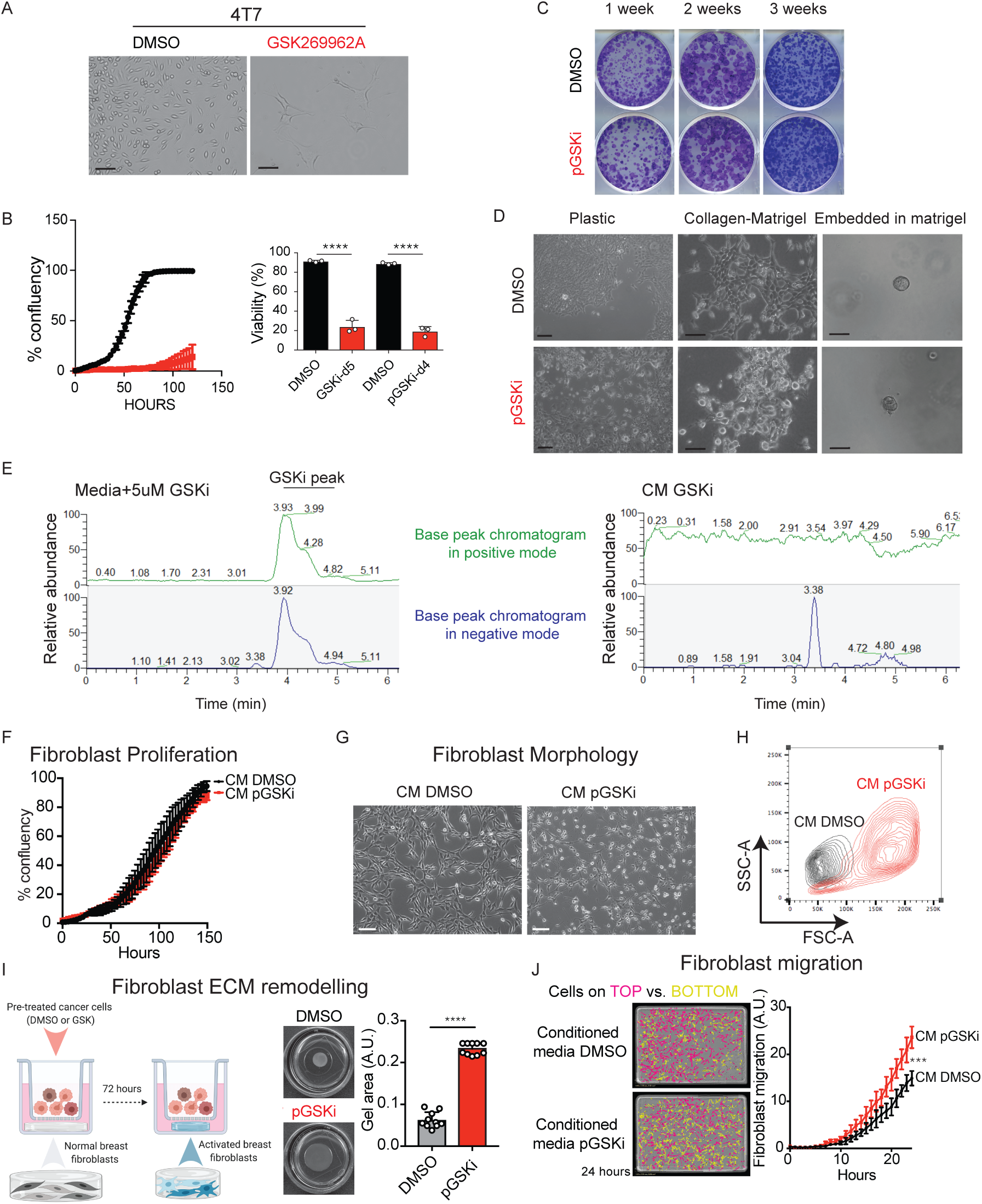
**A.** Representative brightfield images of 4T07 cells following a 5-day treatment with GSKi. **B.** (Left) Growth curve of cells pre-treated with DMSO (black) or GSKi (red). Cell confluency was monitored every 3 hours for <120 hours using Incucyte. (Reft) Percentage of cell viability quantified at 5 days after treatment and at day 4 post-treatment: DMSO (black) or GSKi (red) for 5 days. (Scale bar 100μm). (B-D) Error bars represent the SEM from three independent experiments (dots). Student’s t-test analysis between DMSO and GSKi groups (**** p < 0.0001). **C.** Representative image of the surviving fraction of 4T1 breast cancer cell lines pre-treated with DMSO (black) or GSKi (red) following weekly re-plating and during three consecutive weeks. **D.** Representative bright-field images of pre-treated 4T1 cells with DMSO (upper panels) or GSKi (lower panels) plated on plastic (Left), on a CollagenI-matrigel matrix (Middle) or embedded in matrigel (Right). (Scale bars 50μm). **E.** Liquid chromatography-mass spectrometry chromatograms on positive mode (green) or negative mode (blue) on media containing 5uM of GSKi as positive control (Left) and the 4T1-GSKi pre-treated conditioned-media (Right). The detection limit of 0.01nM was experimentally determined by diluting the positive control. **F.** Growth curve of mNBFs incubated with DMSO (black) or GSKi (red) conditioned media. Cell confluency was monitored every 3 hours for <150 hours using Incucyte. Data were analysed using two-way ANOVA test (ns: not significant). **G.** Representative brightfield images of (mNBF) cell morphology following a 2-day incubation with conditioned media (CM) from DMSO or GSKi pre-treated 4T1 cells. (Scale bar 100μm). **H.** Flow cytometric analysis of the size of mNBFs from (B) based on forward/side scatter. **I.** (Left) Schematic representation of the set-up for fibroblast contraction assays, (middle) representative pictures and (right) quantification of fibroblast contraction assays with DMSO (black) or GSKi (red) pre-treated 4T1 cells. Error bars represent the SEM from three independent experiments and technical repeats (Total n=10 per condition). Student’s t-test analysis (**** p< 0.0001). **J.** (Left) Representative example and (right) quantification of a murine normal fibroblast mNBF chemotaxis migration assay towards DMSO (black) or GSKi conditioned media from 4T1 pre-treated cells. Fibroblast migration data was acquired every hour for 24 hours using Incucyte and the values are given in arbitrary units (A.U.). Data were analysed using two-way ANOVA test (*** p < 0.001).

**Supplementary Figure 2.**
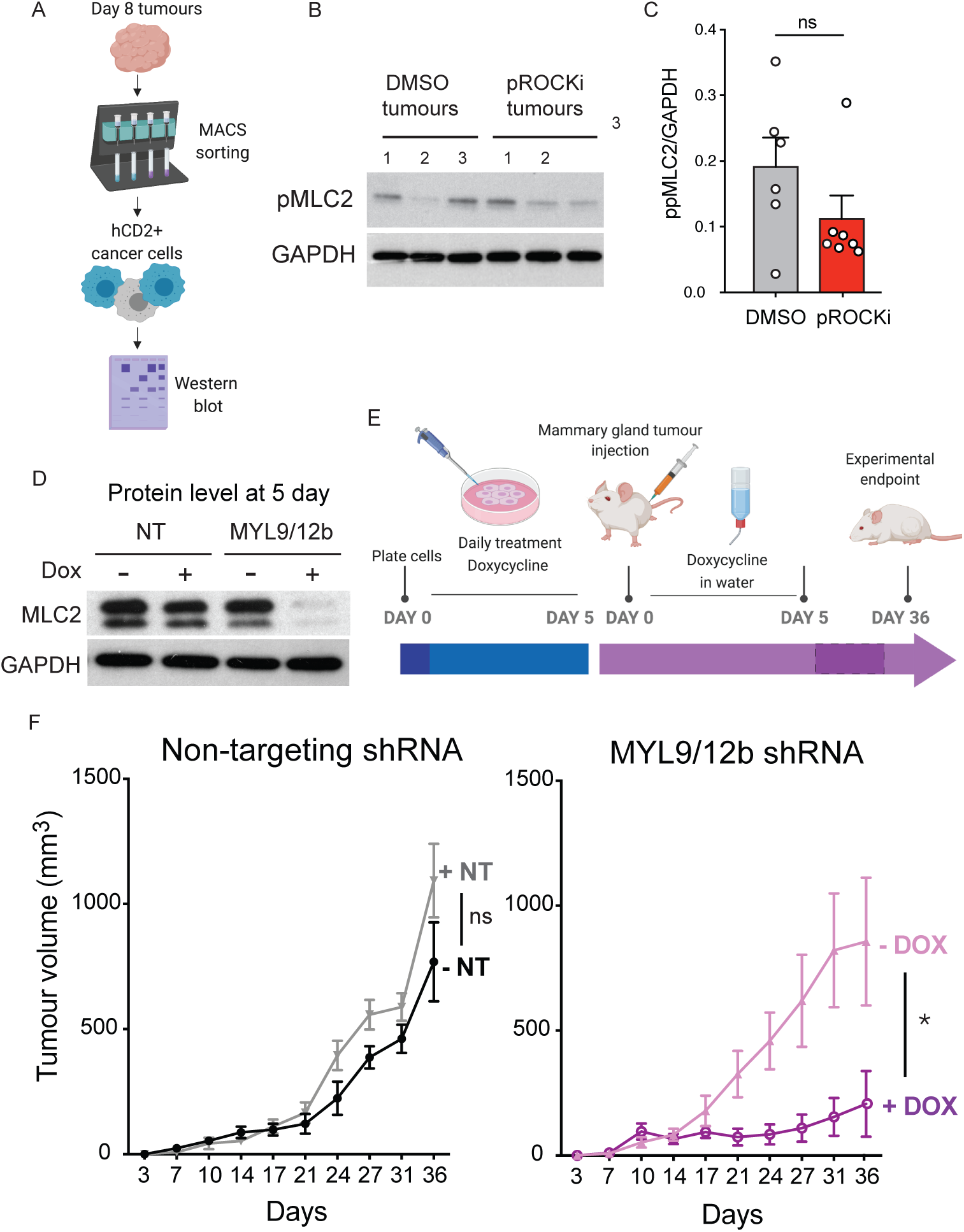
**A.** Schematic of sample collection workflow where hCD2+ cancer cells were isolated from tumours before performing a western blot. **B.** Representative western blot of pMLC2 and GAPDH as loading control in 4T1 pre-treated cells isolated from day 8 tumours. **C.** Quantification of relative ppMLC2 levels in 4T1 tumour-derived cells pre-treated with DMSO (black) or GSKi (red). Error bars represent the SEM from two independent experiments (total n=6-7 tumours). Student’s t-test analysis (ns=non-significant). **D-E.** Western blot analysis of MLC2 after 5 days of DOX treatment *in vitro*. (D). Schematic of the experimental setup: Doxycyline was administered to the cells *in vitro* and *in vivo* during the first five days post-transplantation into the fat pad of Balb/cAnN mice (E). **F.** Mean tumour volume of 4T1 cells transduced with a hairpin inducible vector for Non-targeting (black/grey) or MYL9/12b-targeting shRNA (purple, pink). Error bars represent SEM from two independent experiments (n=8 total number of tumours). Data were analysed using two-way ANOVA with Sidak’s multiple comparisons test. (ns: not significant, * p < 0.05). **I.** Error bars represent the SEM from three independent experiments (total n=17 tumours/condition). Student’s t-test analysis (**** p < 0.0001).

**Supplementary Figure 3.**
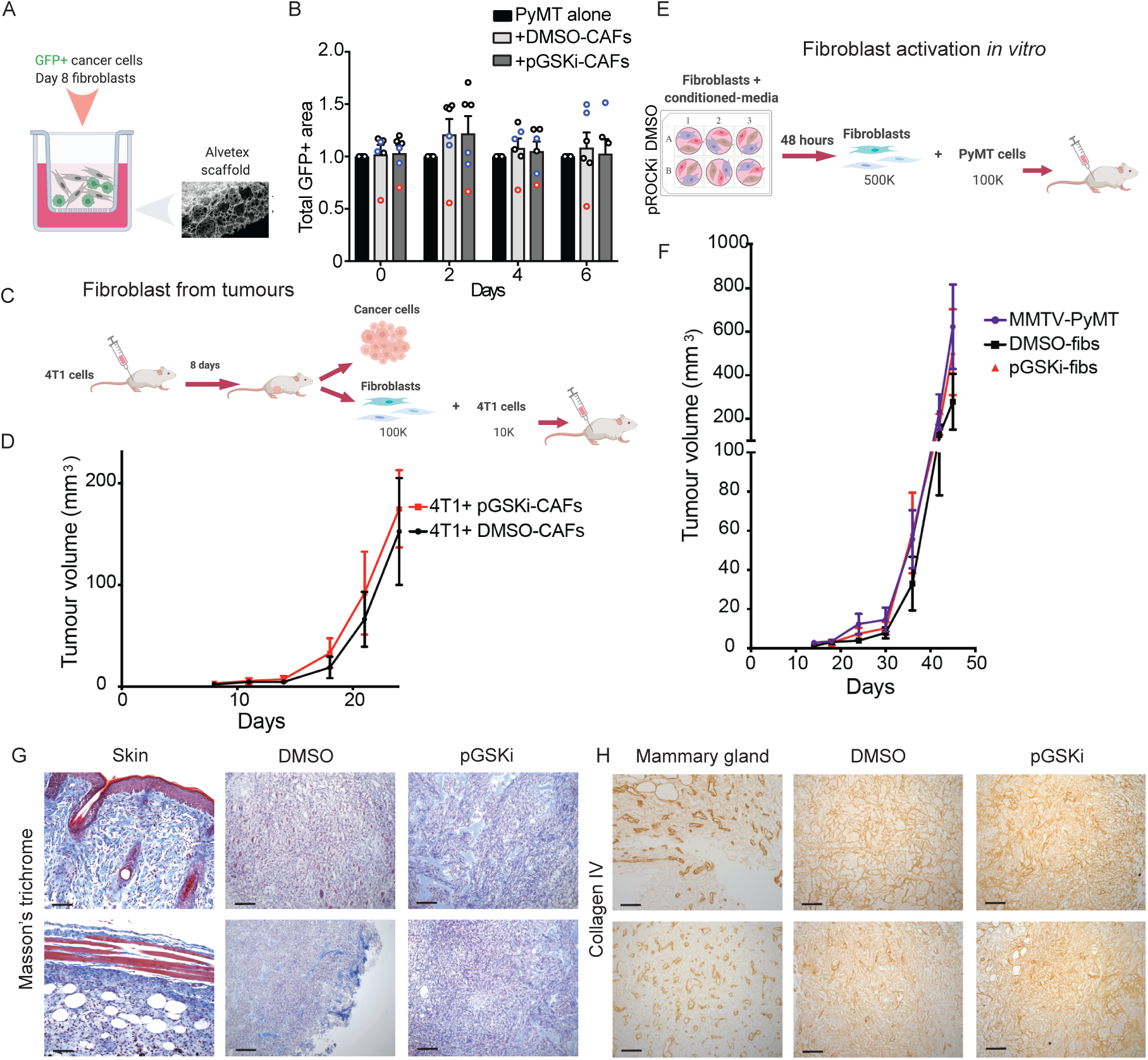
**A.** Experimental design of co-culture on Alvetex scaffolds. **B.** Quantification of MMTV-PyMT-GFP+ cell growth using the total GFP+ area as a readout. Cells were grown alone (black), with DMSO-CAFs (light grey) or GSKi-CAFs (dark grey). Error bars represent the SEM from three independent experiments shown data points. Biological replicates within each experiment are indicated with black, blue or red in the graph. Student’s t-test analysis (ns=non-significant). **C-D.** Diagrams of the experimental design indicating the tumour origin of the CAFs before co-injection with cancer cells (C). Mean tumour volume of 4T1 cells co-injected with DMSO-CAFs (black) or GSKi-CAFs (red) and grown over 24 days (D). Error bars represent the SEM (n=6), Data were analysed using two-way ANOVA (ns=non-significant). **E-F.** Experimental design to generate activated fibroblasts in vitro before co-injection with cancer cells (E). Mean tumour volume of MMTV-PyMT cells alone (purple) or co-injected with DMSO-fibroblasts (black) or GSKi-fibroblasts (red) and grown over 45 days (F). Error bars represent the SEM (n=6). Data were analysed using two-way ANOVA (ns=non-significant). **G.** Representative IHC images of 4T1 tumours 8 days post-injection stained with Masson’s thricrome staining, marking keratin and muscle fibers (red), collagen (blue), and cell nuclei (dark purple). Skin was used as a staining control. **H.** Representative IHC images of 4T1 tumours 8 days post-injection stained collagen IV staining (brown). Normal mammary gland tissue was used as a control.

**Supplementary Figure 4.**
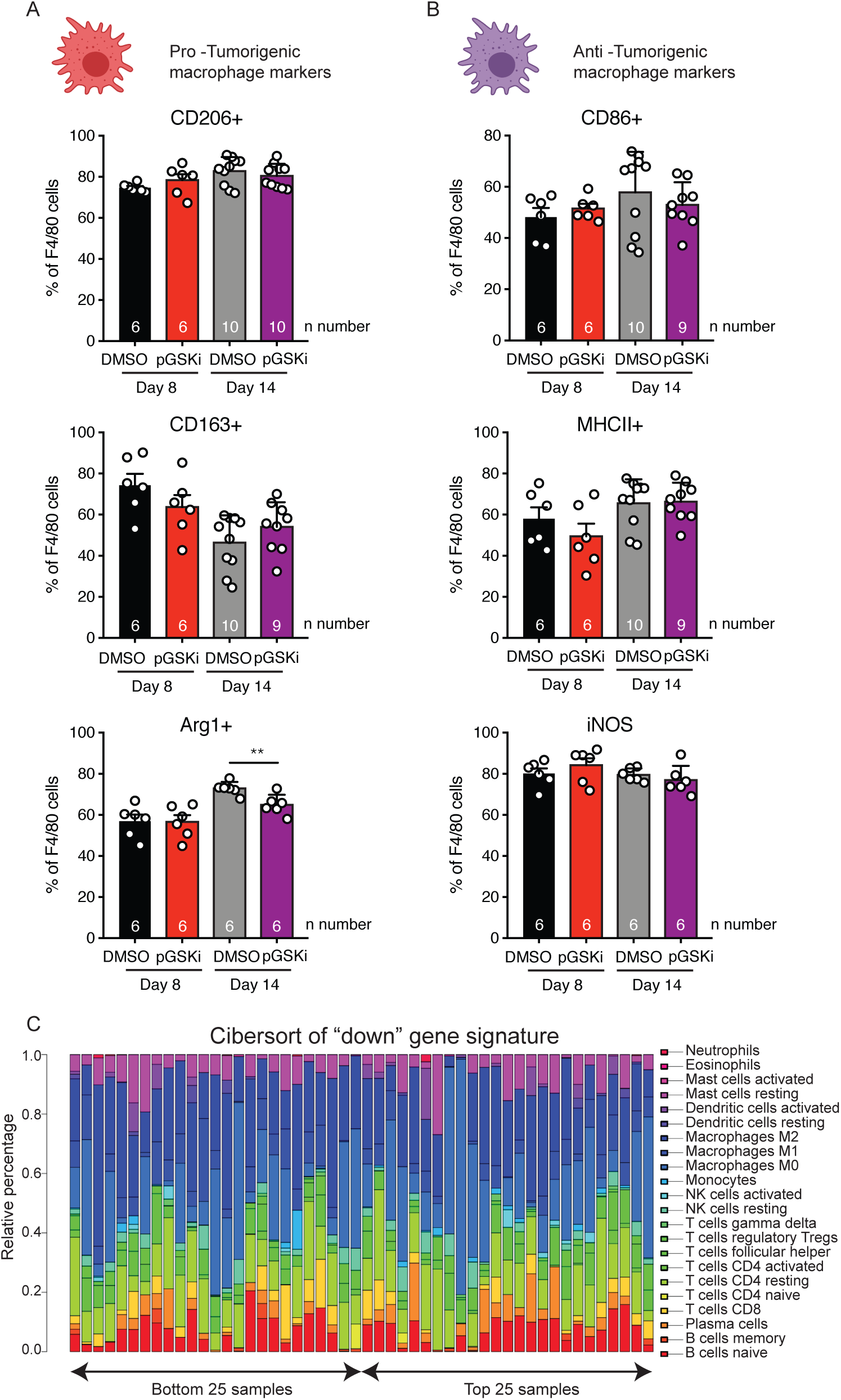
**A-B.** Flow cytometric quantification of pro-tumorigenic (CD206, CD163 and Arg1) (A) and anti-tumorigenic (CD86, MHCII and iNOS) (B) macrophage polarisation markers 8 and 14 days post-transplantation. Cells were gated on alive F4/80+ and error bars represent the SEM from two independent experiments (n numbers indicated under each graph). Data were analysed using t-test (** p < 0.01). **C.** Cibersort analysis of the relative percentage of immune cell populations in the 25 patients with higher (right – top samples) and lower (left - bottom samples) cumulative gene expression for the down fibroblast signature from Figure 3G. Analysis was performed in the TCGA cohort, and the figure legend (right) indicates the cell populations investigated.

**Supplementary Figure 5.**
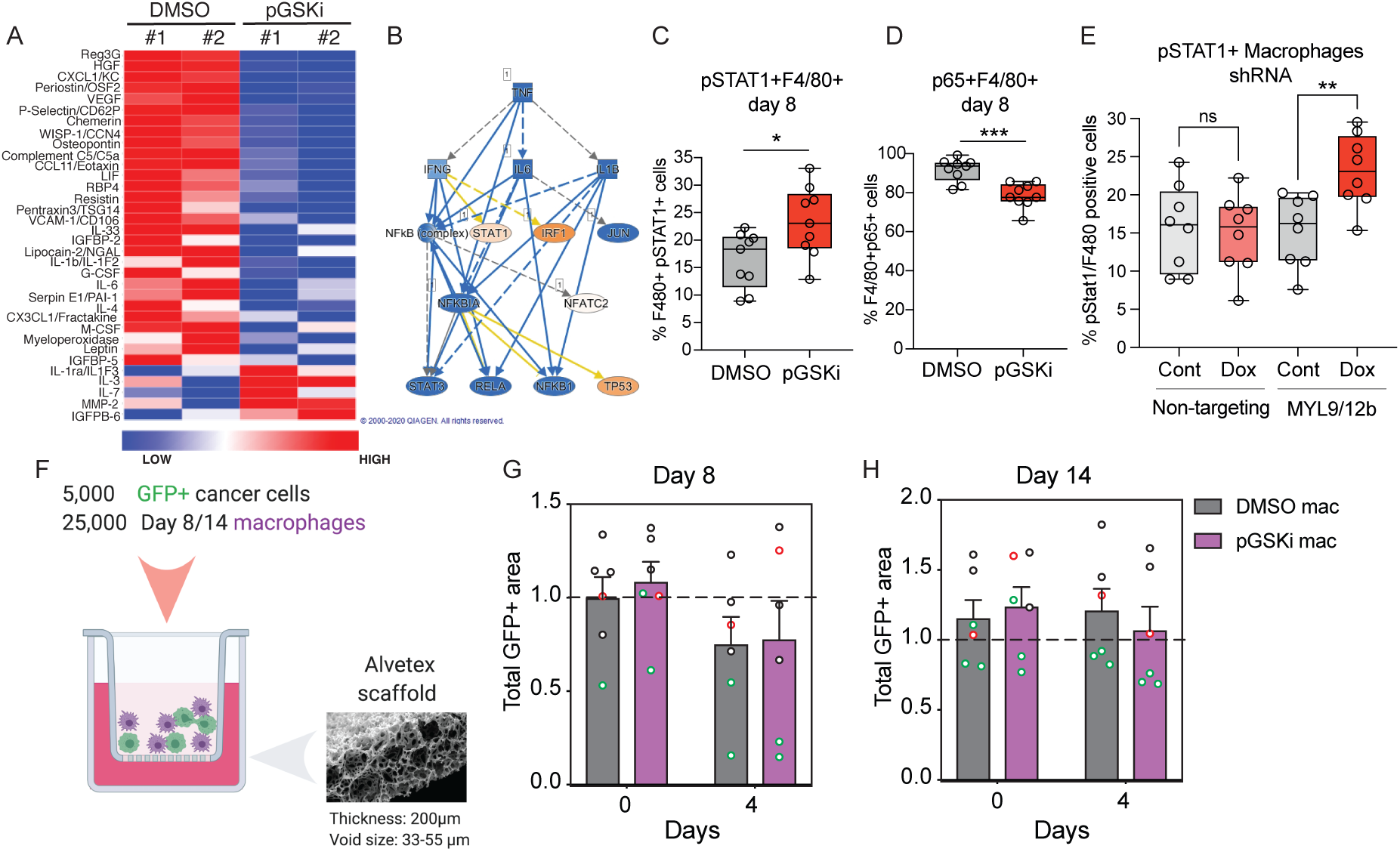
**A.** Heatmap displaying top differentially secreted factors in the fibroblasts from DMSO and GSKi-pre-treated (GSK) tumours at 14 days using comparative marker selection in GennePattern. **B.** Network analysis enriched in GSKi-pre-treated fibroblasts using Ingenuity pathway analysis (RNA signature from Figure 3D). **C.** Percentage of pSTAT1 positive macrophages derived from DMSO (grey) or GSKi (red) pretreated tumours from day 8. **D.** Percentage of p65 positive macrophages derived from DMSO (grey) or GSKi (red) pretreated tumours from day 8. **E.** Percentage of pSTAT1 positive macrophages derived from control or DOX treated of non-targeting or MYL9/12b-targeting shRNA (Fig. 2F-H). **F.** Experimental design of the co-culture. **G-H.** Quantification of MMTV-PyMT-GFP+ cell growth by measuring the total GFP+ area as a readout over the course of 4 days. Cells were grown with DMSO-Macrophages (grey) or GSKi-macrophages (purple) from day 8 (G) or day 14 (H) tumours and normalised to the growth of MMTV-PyMT cells alone (dash line). Error bars represent the +SEM from three independent experiments. Biological replicates within each experiment are indicated with black, green or red data points in the graph. No statistically significant changes were observed between groups following student’s t-test analysis.

## Material and Methods

### Data Availability

The RNaseq datasets generated during this study are available will be made available at the Gene Expression Omnibus and they can currently be provided to the reviewers upon requests.

### Statistical analysis

Unless otherwise stated, statistical analyses were performed using Prism v.7.0c (GraphPad Software). For column bar or scatter dot plots, error bars are the average ± Standard error of the mean (SEM). Boxplots show min to max values or 10-90 percentile. P values were obtained by Student t-tests with paired or unpaired samples when analysing the differences in one variable between two experimental groups. Two-way ANOVA was used where indicated to perform multiple variable analyses or multiple comparisons between experimental groups. Significance was set at P < 0.05 and either the actual P value or symbols describing it (*, p<0.05; **, p<0.01; ***, p<0.001; ****, p< 0.0001) are displayed in graphs.

Data were pooled from at least two independent experiments and the exceptions are specified in the respective figure legends. n values represent biological replicates, with the exception of the 3D Alvetex scaffold assays, for which both technical and biological replicates are shown.

### Bioinformatic analysis

Gene expression data were analysed by the Bioinformatics & Biostatistics facility at the Francis Crick Institute.

Biological replicate libraries were prepared using the polyA KAPA mRNA HyperPrep Kit and sequenced on Illumina HiSeq 4000 platform, generating ∼25 million 76bp single-end reads per sample. Read quality trimming and adaptor removal was carried out using Trimmomatic (version 0.36) (25). The RSEM package (version 1.3.30) (26) in conjunction with the STAR alignment algorithm (version 2.5.2a) (27) was used for the mapping and subsequent gene-level counting of the sequenced reads with respect to Ensembl mouse GRCm.38.89 version transcriptome. All parameters were run as default except “–forward-prob” which was set to 0.5. Normalisation of raw count data and differential expression analysis was performed with the DESeq2 package (version 1.18.1) (28) within the R programming environment (version 3.4.3) (29). Differentially expressed genes were defined as those showing statistically significant differences (FDR <0.05) in the GSKi group relative to the Control Group for both Fibroblast and Macrophage datasets. To determine pathway and biological process enrichment, Clarivate Analytics’s Metacore pathway analysis tools was used to determine pathway and biological process enrichment using a hypergeometric test. [https://portal.genego.com/]. Gene lists ranked by the Wald statistic were used to look for pathway and selected genesets using the Broad’s GSEA software (version 2.1.0) with genesets from MSigDB (version 6) (30).

Differentially regulated genes (logFC > 1 and FDR < 0.05) from the fibroblast data analysis were converted to human orthologs using HomoloGene [https://www.ncbi.nlm.nih.gov/homologene]. Breast Carcinoma (BRCA) normalised expression data was downloaded from the FireHose Broad GDAC website [https://gdac.broadinstitute.org/] Data downloads were all version 2016012800.0.0. BRCA patients were ranked using the sum of expression values of down regulated gene signature. The top quartile of patients were names as “HIGH”, and the bottom quartile named “LOW”. Survival plots were generated using R’s survival package (version 2.41-3) (31) and the level of statistical significance determined using the log rank test.

### Animal procedures

Mice used were: Wildtype mice in BALB/CJ background, MMTV-PyMT+, MMTV-PyMT+ actin GFP mice and Rag1-/- in pure genetic FVB/N background and.

Animals were maintained under specific pathogen-free conditions and handled in accordance with the Institutional Committees on Animal Welfare of the UK Home Office and all the procedure were performed under the PPL P83B37B3C licence.

### Cell lines

This study used a variety of breast cancer cell lines derived from the mouse normal breast fibroblasts (mNBF) were a gift from E. Sahai and MMTV–PyMT cells were isolated from MMTV–PyMT tumours (FVB/N background) as previously described (4). The 4T1 and 4T07cell lines were provided by the Cell Services Unit of The Francis Crick Institute.

4T1 cells plated in 2D were treated with either vehicle (DMSO) or 5 μM of GSK269962A (Axon, 1167) for five days before performing in vitro and in vivo experiments, unless stated differently. Media and inhibitors were changed daily. Matrices of 2:1 Pure Collagen-I and Matrigel were prepared to a final collagen concentration of ∼4 mg/ml and a Matrigel concentration of ∼2mg/ml. The cells were seeded on top in 200ul of growth medium.

### Generation of shRNA plasmids

The doxycycline inducible Tet-PLKO-puro lentiviral plasmid vector used in this study was purchased from Addgene (#21915, #21916). The vector contains all the necessary components for the controlled doxycycline-triggered expression of shRNA in mammalian cells. Gene knock-down is triggered by the presence of doxycycline in the growth media, while in the absence of doxycycline, shRNA expression is blocked by the presence of constitutively-expressed TetR protein.

Forward and reverse oligonucleotides for a MYL9 and MYL12b-targeting shRNA and a nontargeting shRNA construct were designed using The RNAi Consortium collection (MISSION® shRNA, Sigma Aldrich).

### MACS isolation of cell subsets from tumour

Tumour tissues were dissociated and processed to single-cells as described previously (4). MidiMACS separators and LD and LS columns (Milteny, 130-042-901, 130-042-401) were used to magnetically isolate mouse cellular subsets from tumour tissue at day 8 and day 14 and the procedure was performed on ice and promptly.

For fibroblast isolation, cells were isolated from Balb/cAnN mice orthotopically transplanted with 4T1 tumours pre-treated with DMSO or GSK. Single-cell suspensions were incubated with mouse FcR Blocking Reagent and subsequently with the following APC labelled antibodies – CD45, CD31, CD326 and hCD2. Cells were washed and incubated with magnetic anti-APC MicroBeads (Miltenyi, 130-090-855). Magnetically labelled immune, endothelial, epithelial and cancer cells were isolated using LD columns, washed thrice with 3 ml of MACS and the unlabelled flow-through cell fraction was the mesenchymal cells that we will refer to as fibroblasts hereafter. To increase the purity of the cells, the eluted fraction was applied to a second round of separation with a new LD column.

For macrophage isolation, F4/80+ cells were obtained by using the Anti-F4/80 MicroBeads UltraPure kit (Milteny, 130-110-443) following the manufacturer’s instructions. Approximately 2×10^7^ single-cells from the tumour were stained with a mix of 180 μl MACS buffer and 20 μl of Anti-F4/80 MicroBeads for 15 minutes at 4°C in the dark. During this incubation, an LS column was placed onto a MidiMACS separator where magnetic field was applied, and the column was rinsed with 3 ml of MACS buffer. Cells were washed twice with 4 ml of MACS buffer and resuspended in 1ml prior to proceeding to magnetic separation by applying the cell suspension onto the column. Columns were washed thrice with 3 ml of MACS buffer before being carefully removed from the separator and placed on a FACS tube. 4.5 ml of MACS buffer was added onto the LS column and the plunger was firmly pushed to elute the F4/80+ cell fraction which had been retained within the column due to the F4/80+ positive selection. To increase the purity of the cells, the eluted fraction was applied to a second round of separation with a new LS column.

### Alvetex scaffold 3D assay

Alvetex Scaffold 96-well plates (ReproCELL, AVP009) were washed once with 70% ethanol to render it hydrophilic. Following three washes with PBS, they were coated with a collagen-solution and incubated for 1 hour at 37°C prior to cell seeding. Primary MMTV–PyMT actin–GFP cells were seeded at a density of 5,000 cells per well and fibroblasts and macrophages isolated from tumours were seeded at a density of 25,000 cells per well in a total volume of 100ul of complete MMTV-PyMT media.

The growth of GFP+ cells was monitored every two days using the SteREO LumarV12 stereomicroscope (Zeiss) and for a total of 6 days. Images were taken at a magnification of 19.4X, Z +0.09, exposed for 2 seconds and quantified using ImageJ software. For quantification, images were stacked and the Li’s minimum cross entropy thresholding algorithm was used. All data was normalized to the growth of PyMT-GFP cells at each timepoints.

### Generation of conditioned media

100,000 4T1 cells (control or GSKi treated) were plated on tissue culture treated 24-well plates in 1 ml of complete DMEM media, which was conditioned at 37°C for 48 h. The medium preparation was collected and spun at 1200 rpm for 20 min. The supernatant after this centrifugation was collected and used as conditioned media (CM).

### Fibroblast contraction assay

To assess fibroblast activation via their ability to remodel an extracellular matrix, 7,5×10_4_ mNBF cells were embedded in 100 μl of 2:1 collagen-I:Matrigel and seeded on a 35-mm glass-bottom dish (MatTek, P35-1.5-14-C). Once the gel had polymerized, a low height co-culture insert (Millipore, PICM-ORG-50) was placed on the dish and 1,5×10_5_ tumour cells were seeded on the insert in 1ml of complete DMEM media. Gel contraction was monitored daily by taking photographs of the gels. The gel contraction value refers to the contraction observed after 3 days, which was measured using ImageJ software and calculated as the gel area in arbitrary units (A.U.).

### Cell proliferation assays – incucyte

Cells were plated sub-confluently with numbers ranging from 1,000 to 10,000 cells per well into 96-well high content imaging black plates (Corning, CLS4580). Cell growth was monitored over 4-6 days using IncuCyteZOOM® Live Cell Imaging. All conditions were performed in triplicate and 4 images per well were acquired every 3 hours. The total area covered by cells was automatically measured by the software through confluency masks. The percentage of confluency over time was calculated as the average area covered by cells relative to the total well area.

### Cell migration assays

Chemotaxis of mNBF fibroblasts was assessed using Incucyte Clearview 96-well chemotaxis plate with ,2% pore density following the manufacturer’s protocol (Essen Bioscience, 4582). 1,000 mNBF cells in DMEM with 1% FBS were loaded in the upper chamber compartment. CM from 4T1 DMSO or GSK was loaded in the lower chamber. Migration was monitored using the IncuCyteZOOM® Live Cell Imaging, and images of the upper and lower chambers were taken every hour. The total area covered by cells was automatically measured by image segmentation using the integrated chemotaxis analysis plugin.

### Flow cytometry (FACS)

Prepared single-cell suspensions of mouse tissues and in vitro treated cancer cells were incubated with mouse FcR Blocking Reagent (Miltenyi, 130-092-575) for 10 min at 4°C followed by a 30 minute incubation at 4°C in the dark with a combination of the pre-labelled antibodies. Following two washes with MACS buffer, dead cells were stained with DAPI (4,6-Diamidino-2-phenylindole dihydrochloride, Sigma, D9542) and analysed with an LSRFortessa cell analyser running FACSDiva software (BD Biosciences) and FlowJo 10.4.2 software (FlowJO, LCC 2006-2018). All cell-sorting experiments were carried out using a BD Influx cell sorter (BD Biosciences).

### Antibodies

Details of the primary antibodies used for immunohistochemistry in this study are listed below.

Immunofluorescence

| Antigen | Source Ig | Reactivity against | Dilution | Antigen retrieval | Catalogue number |
| --- | --- | --- | --- | --- | --- |
| SMA | Mouse | Mouse, Human | 1:750 | Citrate | DAKO, M0851 |
| Endomucin | Rat | Mouse |  | Citrate | sc-65495 |
| Collagen IV | Rabbit | Mouse | 1:500 | Citrate | USBiological C7510-50B |
| F4/80 | Rat | Mouse | 1:100 | Trypsin | eBioscience, 14-4801 |
| pSer19-MLC2 | Rabbit polyclonal | Mouse, Human | 1:50 | Access super buffer | CST, 3671L |

FACS

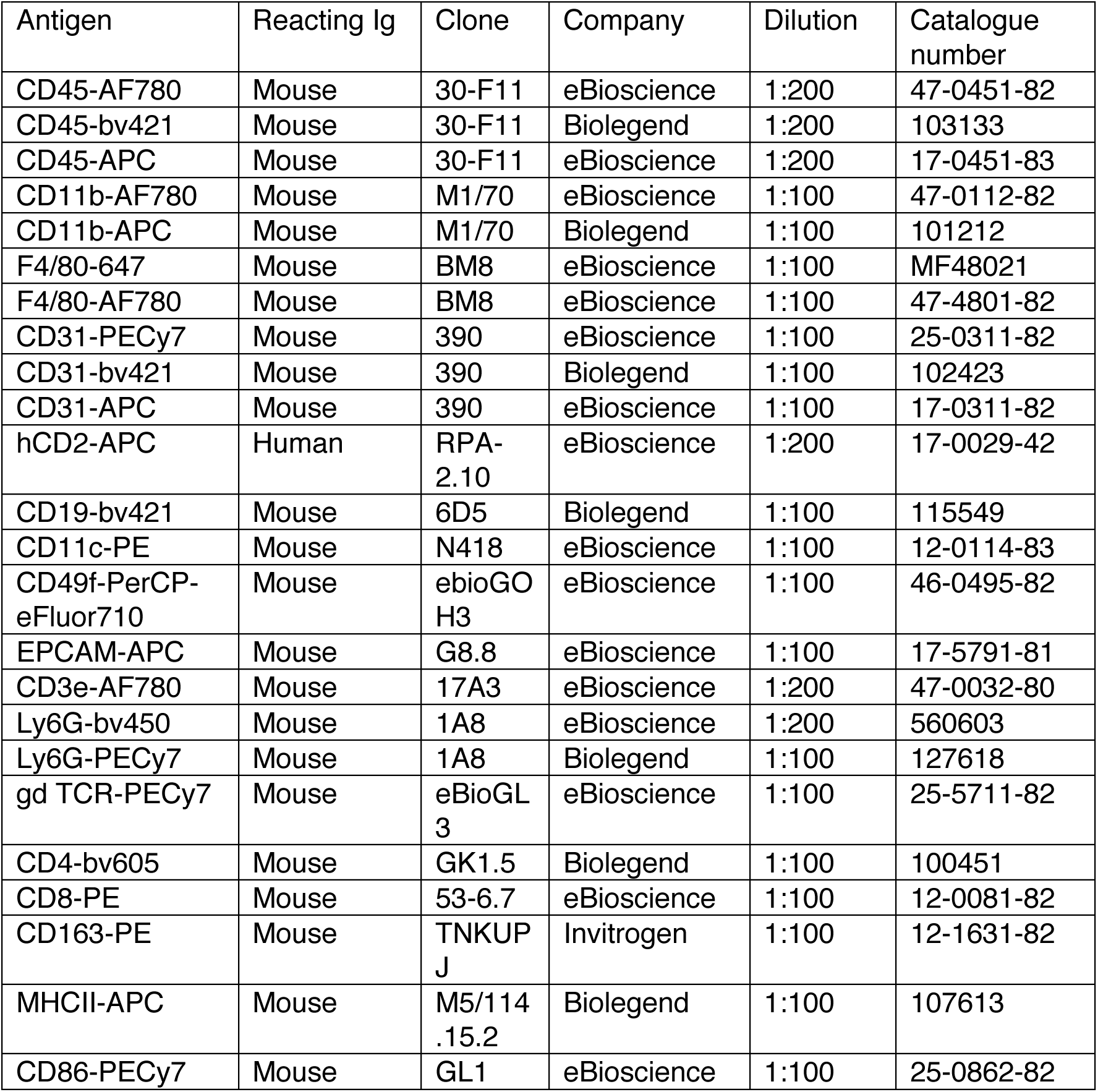
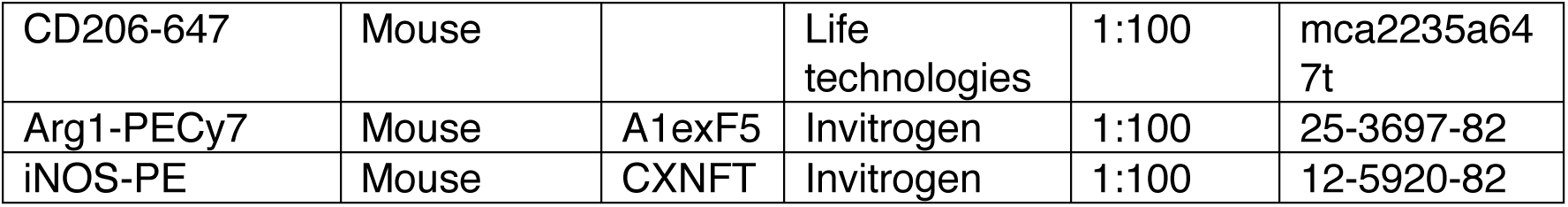

| Antigen | Reacting Ig | Clone | Company | Dilution | Catalogue number |
| --- | --- | --- | --- | --- | --- |
| CD45-AF780 | Mouse | 30-F11 | eBioscience | 1:200 | 47-0451-82 |
| CD45-bv421 | Mouse | 30-F11 | Biolegend | 1:200 | 103133 |
| CD45-APC | Mouse | 30-F11 | eBioscience | 1:200 | 17-0451-83 |
| CD11b-AF780 | Mouse | M1/70 | eBioscience | 1:100 | 47-0112-82 |
| CD11b-APC | Mouse | M1/70 | Biolegend | 1:100 | 101212 |
| F4/80-647 | Mouse | BM8 | eBioscience | 1:100 | MF48021 |
| F4/80-AF780 | Mouse | BM8 | eBioscience | 1:100 | 47-4801-82 |
| CD31-PECy7 | Mouse | 390 | eBioscience | 1:100 | 25-0311-82 |
| CD31-bv421 | Mouse | 390 | Biolegend | 1:100 | 102423 |
| CD31-APC | Mouse | 390 | eBioscience | 1:100 | 17-0311-82 |
| hCD2-APC | Human | RPA-2.10 | eBioscience | 1:200 | 17-0029-42 |
| CD19-bv421 | Mouse | 6D5 | Biolegend | 1:100 | 115549 |
| CD11c-PE | Mouse | N418 | eBioscience | 1:100 | 12-0114-83 |
| CD49f-PerCP-eFluor710 | Mouse | ebioGO H3 | eBioscience | 1:100 | 46-0495-82 |
| EPCAM-APC | Mouse | G8.8 | eBioscience | 1:100 | 17-5791-81 |
| CD3e-AF780 | Mouse | 17A3 | eBioscience | 1:200 | 47-0032-80 |
| Ly6G-bv450 | Mouse | 1A8 | eBioscience | 1:200 | 560603 |
| Ly6G-PECy7 | Mouse | 1A8 | Biolegend | 1:100 | 127618 |
| gd TCR-PECy7 | Mouse | eBioGL 3 | eBioscience | 1:100 | 25-5711-82 |
| CD4-bv605 | Mouse | GK1.5 | Biolegend | 1:100 | 100451 |
| CD8-PE | Mouse | 53-6.7 | eBioscience | 1:100 | 12-0081-82 |
| CD163-PE | Mouse | TNKUP J | Invitrogen | 1:100 | 12-1631-82 |
| MHCII-APC | Mouse | M5/114 .15.2 | Biolegend | 1:100 | 107613 |
| CD86-PECy7 | Mouse | GL1 | eBioscience | 1:100 | 25-0862-82 |
| CD206-647 | Mouse |  | Life technologies | 1:100 | mca2235a64 7t |
| Arg1-PECy7 | Mouse | A1exF5 | Invitrogen | 1:100 | 25-3697-82 |
| iNOS-PE | Mouse | CXNFT | Invitrogen | 1:100 | 12-5920-82 |

